# Lysosomal escape and TMEM106B fibrillar core determine TDP-43 seeding outcomes

**DOI:** 10.64898/2025.12.19.695531

**Authors:** Weijia Zhong, Carlo Scialò, Beatrice Gatta, Manon Häfliger, Noemi Leu, Flavio Lurati, Martina Peter, Nandini Ramesh, Bernd Roschitzki, Somanath Jagannath, Ruchi Manglunia, Elena de Cecco, Su Min Lim, Oscar G Wilkins, Adriano Aguzzi, Michael Ward, Pietro Fratta, Leonard Petrucelli, Clotilde Lagier-Tourenne, Magdalini Polymenidou

**Author notes:** These authors contributed equally to this work (Weijia Zhong, Carlo Scialò).

## Abstract

Frontotemporal lobar degeneration with TDP-43 inclusions (FTLD-TDP) shows striking clinical and neuropathological heterogeneity, yet a systematic analysis of subtype-specific features and inter-patient variability was missing. We treated human neurons and neuron-like cells with 30 postmortem brain samples and quantified neoaggregate formation, loss of function and changes in the TDP-43 interactome to define determinants of seeding outcomes. Potent FTLD-TDP-A seeds drove a progressive collapse of physiological TDP-43 interactions accompanied by functional loss. Beyond the burden of pathological TDP-43, we identified the fibrillar core of the lysosomal protein TMEM106B as a critical pro-seeding factor. Transient lysosomal injury markedly enhanced neoaggregation and loss of function, likely by promoting fibril interactions with native TDP-43. Our work establishes a mechanistic link between TMEM106B and TDP-43 aggregation, identifies lysosomal escape as a key driver of pathology and introduces the strongest model yet for seeded TDP-43 aggregation and loss of function, to enable discovery of disease modifiers.

## Introduction

TAR DNA-binding protein-43 (TDP-43) cytoplasmic aggregation and depletion from its physiological nuclear localization are the pathological hallmarks of frontotemporal lobar degeneration with TDP-43 inclusions (FTLD-TDP) and nearly all cases of amyotrophic lateral sclerosis (ALS)^1–3^. This dual pathology drives neuronal loss through a combination of gain-of-toxicity and loss-of-function mechanisms^4–7^. Reduced nuclear TDP-43 leads to failure in governing key RNA processing events^8–13^, ultimately causing the misregulation of critical neuronal proteins, including *STMN2*^7, 14^, *UNC13A*^4, 15^, *KCNQ2*^16^ and *NPTX2*^17^.

Despite the unifying TDP-43 pathology across ALS and FTLD-TDP patients, its pathological aggregates exhibit remarkable variability in histological patterns, and morphological, structural, and biochemical properties^18–22^. Among those, FTLD-TDP type A (FTLD-TDP-A) and type C (FTLD-TDP-C) represent particularly divergent pathological entities^23–25^. FTLD-TDP-A is characterized by abundant neuronal cytoplasmic inclusions and dystrophic neurites in superficial cortical laminae, typically associated with progranulin mutations and prominent clinical apathy and social withdrawal. In contrast, FTLD-TDP-C displays axonal pathology^5^ primarily in superficial cortical layers, is predominantly sporadic, and patients with this pathology have slower disease progression and older age at death compared to those with other FTLD-TDP subtypes^18, 26^. Notably, the TDP-43 fibrils of the two subtypes were recently shown to have distinct structures^27, 28^ with pathological FTLD-TDP-C bearing unique heterotypic amyloid filaments comprising TDP-43 co-assembled with another protein, annexin A11 (ANXA11)^28^, a finding supported by neuropathological^29^ and genetic^30, 31^ observations. Furthermore, the type II transmembrane lysosomal protein TMEM106B was found to form amyloid filaments in aging brains and across a range of neurodegenerative diseases^32–34^ while genetic variants of TMEM106B determine disease progression rates of FTLD patients, with protective single nucleotide polymorphisms (SNPs) associated with longer survival after disease onset^35–37^. The clinical manifestations of ALS and FTLD exhibit spatiotemporal progression patterns, compatible with a prion-like propagation mechanism^38–41^. Since our initial proposal^42^ evidence supporting non-cell-autonomous spread of TDP-43 pathology as a key pathological mechanism in ALS and FTLD has accumulated^5, 18–22, 43–46^. We showed that addition of patient-derived pathological TDP-43 to cultured cells is directly neurotoxic^18^ and induces the templated aggregation of intracellular TDP-43^5^ in a disease subtype-specific fashion, as evidenced by the size and structure of neoaggregates, the potency of seeding and the severity of neurodegeneration, with the latter two correlating with the rate of disease progression in patients. In line with our data, seeding with TDP-43 extracts derived from brains of patients with different FTLD subtypes also produced distinctive patterns of TDP-43 pathology *in vivo*^19, 20^. More recently, we exploited the templating properties of TDP-43 fibrils to establish a cellular model that unambiguously links TDP-43 aggregation to loss of function and circumvents the need for overexpression or knockdown^44^. We showed that *in vitro* generated TDP-43 fibrils were abundantly internalized by human neuron-like cells, efficiently recruited endogenous TDP-43, and formed cytoplasmic inclusions reminiscent of ALS/FTLD pathology. Combining a fluorescent reporter of TDP-43 function (TDP-REG^47^) with RNA sequencing and proteomics, we demonstrated a loss-of-function profile resulting from TDP-43-templated aggregation^44^.

Here, we exploited this cellular seeding model to dissect the molecular determinants of TDP-43 seeding outcomes and elucidate the contribution of its proteoforms and other insoluble components to the induction of neoaggregation and subsequent loss-of-function. To identify changes in the interactome of endogenous TDP-43 during aggregation, we integrated the proximity-dependent biotin identification (BioID)^48^ approach with our seeding model in neuron-like SH-SY5Y cells. Our proteomic analysis revealed changes in TDP-43 interactors following exposure to pathological TDP-43 extracted from FTLD-TDP-A and FTLD-TDP-C postmortem brains, including proteins involved in the protein-nucleic acid complex assembly as a shared significantly decreased pathway, pointing to loss of physiological nuclear TDP-43 interactions. Moreover, we identified increased binding of TDP-43 to actin-associated proteins, upon treatment with patient brain extracts initially from both FTLD subtypes but persisting only in the FTLD-TDP-A potent seeders. Using a cohort of 30 postmortem samples, we quantified neoaggregate formation and loss of function over time. Importantly, the population of cells harboring detectable exogenous TDP-43 aggregates declined over time – consistent with degradation of the internalized fibrils – while cells exhibiting induced loss of function increased over the same period, indicating a propagating pathological cascade.

To identify the factors shaping the seeding outcomes, we quantified biochemical signatures in patient-derived seeds and assessed how these features related to neoaggregate formation and TDP-43 loss of function. Our analyses highlighted phosphorylated TDP-43 and TMEM106B levels as the strongest predictors of neoaggregate formation. Unexpectedly, TMEM106B alone emerged as the sole feature associated with loss of function—an observation we independently validated in a new neuronal model incorporating the TDP-REG. To explore whether lysosomal integrity contributed to these effects, we transiently disrupted lysosomal membranes after exposing cells to exogenous TDP-43 aggregates and observed a marked enhancement of the resulting loss of function. Collectively, our findings identify the TMEM106B fibril core within patient-derived seeds as the principal driver of aggregation-linked TDP-43 dysfunction and reveal lysosomal escape as a previously unrecognized mechanism that accelerates prion-like TDP-43 pathology.

## Results

### Proximity proteomics identify nuclear and cytoplasmic interactors of TDP-43

To capture the changes in the protein interactors of TDP-43 under normal and pathological conditions, we adapted the BioID method^48^, which is based on the fusion of a promiscuous mutant of the *A. aeolicus* biotin ligase (BioID2) to a bait protein to catalyze the biotinylation of proximate proteins *in situ* within the natural cellular environment^49^ (**Fig. 1a**). We generated stable SH-SY5Y cells expressing BioID2 fused to either wild type TDP-43 (WT), or a variant carrying mutations in the nuclear localization signal to abolish active nuclear import (mNLS)^50,51^. To distinguish true TDP-43 interactors from non-specific biotinylating events, we generated a third line expressing BioID2 fused to GFP (**Fig 1b**). After cell lysis, biotinylated interactors were captured by streptavidin-coated beads, followed by mass spectrometry analyses. When our stable lines were incubated with excess biotin in culture, proximity-dependent biotinylation of BioID2-TDP-43WT- and BioID2-TDP-43mNLS-associated proteins showed distinct patterns on Western blots (**Fig. 1c**). The lower expression of BioID2-TDP-43mNLS, compared to its WT counterpart may be due to toxic effects of this cytoplasmic TDP-43 variant. Importantly, biotinylated interactors followed the subcellular localization of the bait proteins, i.e. were predominantly found in the nucleus of BioID2-TDP-43WT-expressing cells, or both cytoplasmic and nuclear compartments in BioID2-TDP-43mNLS-expressing cells (**Fig. 1d**). Principal component analysis (PCA) of the enriched biotinylated proteome demonstrated a clear separation of the three conditions (**Fig. 1e**). 713 proteins were identified specifically interacting with BioID2-TDP-43WT (**Fig. 1f**) and 473 proteins with BioID2-TDP-43mNLS (**Fig. 1g**). To identify functional clusters among the interactors, we performed Gene Ontology (GO) enrichment analysis. Wild type TDP-43-specific interactors were strongly enriched in splicing factors, consistent with the well-known and prominent role of TDP-43 in RNA splicing and in line with previous interactome studies (**Fig. 1h**; **Table S1, S2**)^52–61^. In addition, our analysis uncovered previously known TDP-43 molecular functions, such as transcription regulation and DNA binding (**Fig. S1**)^3, 11, 12, 62^. Validating our approach, the differences in the pathway-specific enrichment and interaction networks between TDP-43WT and TDP-43mNLS proteomes (**Fig. S1**), pointed to a shift from nuclear interactomes in the wild type (**Fig. 1h**) to cytoplasmic ones in the mutant NLS variant, which were enriched in translation regulation, RNA catabolic processes and the assembly of membraneless organelles, such as stress granules (**Fig. 1i**).

**Figure 1.**
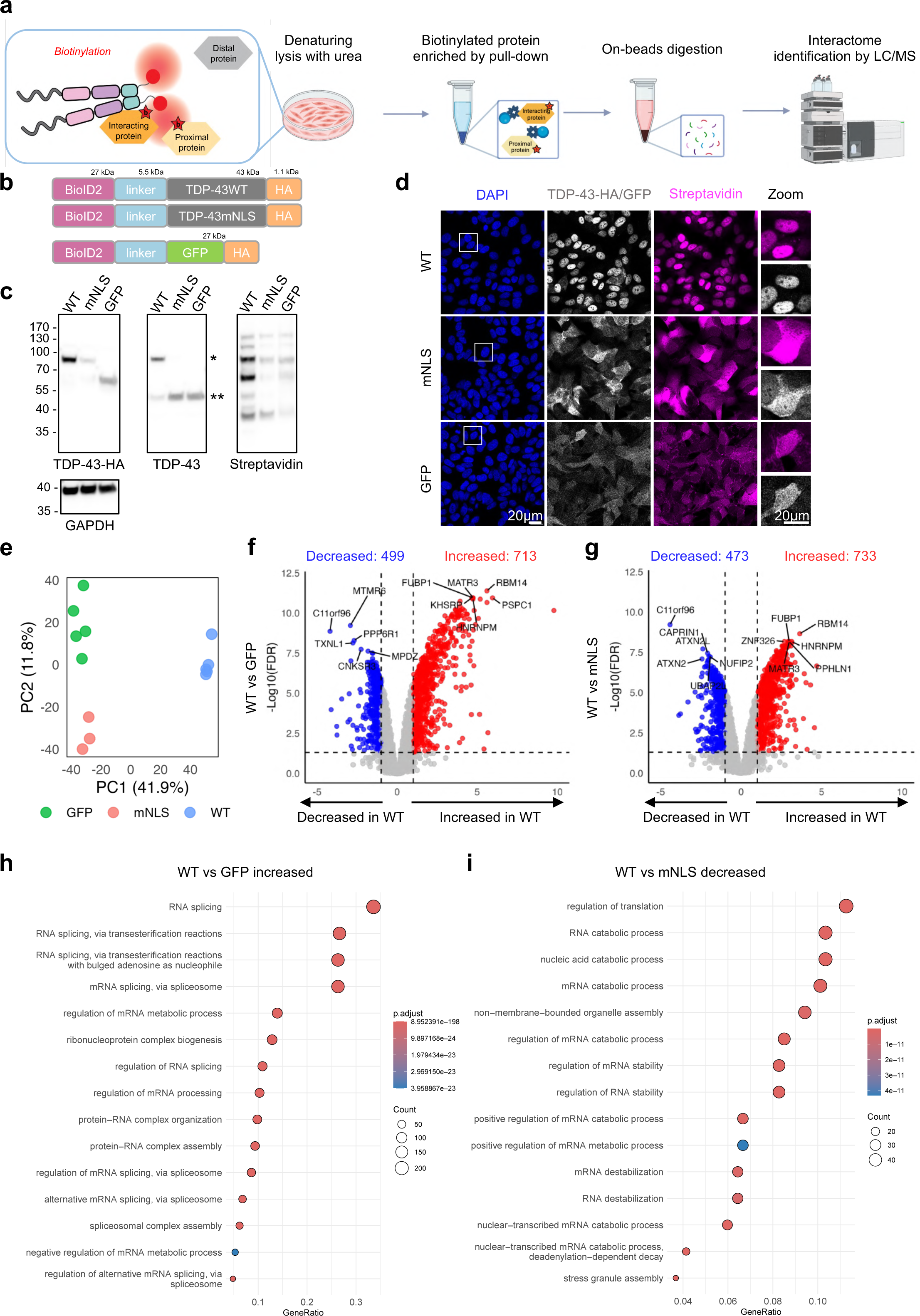
Proximity proteomics identify nuclear and cytoplasmic interactors of TDP-43. **(a)** Schematic representation for the proximity-dependent biotinylation and identification (BioID) workflow for the interactors of TDP-43. (**b**) Domain structures of BioID2-TDP-43 WT/mNLS constructs. (**c**) Western blot with an anti-HA (left), anti-TDP-43 (middle) and streptavidin-HRP (right) antibodies, showing the presence of transgene (*) and endogenous (**) TDP-43. GAPDH is shown as a loading control (lower panel). (**d**) Representative images depicting the subcellular localization of BioID2-TDP-43 WT/mNLS/GFP. Scale bar: 20 μm. (**e**) Principal component analysis (PCA) demonstrating separation of the three experimental conditions at proteomics level. Each dot represents a biological replicate, *n* = 3. Volcano plots from the comparison of WT vs. GFP (**f**) and WT vs. mNLS (**g**). The dot colors indicate if the interaction of TDP-43 with a protein partner is decreased (blue), increased (red) or not significantly modified (grey) according to the group difference (x-axis) with a threshold of 1-fold and -Log_10_ false discovery rate (FDR) (y-axis) < 0.05. Displayed numbers are the numbers of significantly modified interactions within each comparison. (**h**) Enriched biological processes are shown for increased interaction in the comparison of WT vs. GFP and (**i**) decreased for WT vs. mNLS. Dot sizes represent the number of proteins in each pathway (count), while dot colors indicate pathway significance based on adjusted p-values (p.adjust). The GeneRatio on the x-axis represents the proportion of DEGs associated with each pathway.

### Patient-derived seeds exhibit variable pathological burden and seeding potency

We previously showed that TDP-43 pathological aggregates could be isolated from postmortem brain through nuclease digestion combined with harsh solubilization and a single centrifugation step, a method we termed SarkoSpin^63^. We performed SarkoSpin on brain samples from a cohort comprising 10 individuals with FTLD-TDP-A, 10 with FTLD-TDP-C and 10 controls without neurodegenerative conditions (CTRL). To characterize the pathological protein burden in each sample, we performed immunoblotting and quantified the total (**Fig. 2a-b**), or phosphorylated TDP-43 at positions 409/410 (pTDP-43) (**Fig. 2c, d, Fig. S2a**). We then exposed our SH-SY5Y BioID2-TDP-43WT line (**Fig. 1**) with patient-derived aggregates to test their seeding propensities (**Fig. 2e**). To track cellular uptake, patient-derived SarkoSpin extracts were labeled with the fluorescent dye Atto-488, as previously done^44^. Cells were growth-arrested with biotin depletion^64^ to minimize dilution on the seeding effect caused by cell proliferation. Following trypsinization to eliminate non-internalized aggregates, cells were analyzed at 6, 10 and 18 days post incubation (d.p.i.) by fluorescence-activated cell sorting (FACS) to evaluate uptake of the exogenous aggregates (**Fig. 2e**). All exogenous seeds, regardless of the source exhibited high internalization rate at 6 d.p.i, maintained similar levels at 10 d.p.i., and then showed a marked decline afterwards when assessed at 18 d.p.i. (**Fig. 2f, Fig. S2b**). Despite some individual variability per condition (**Fig. S2c**), the FTLD patient seeds displayed an overall higher rate of retention within cells compared to the extracts from the non-neurodegenerative control brains, with the FTLD-TDP-A seeds showing the highest cell retention (**Fig. 2f**). The internalization rate was largely unchanged in our BioID-TDP-43mNLS cell line with a subset of patient-seeds (**Fig. S2d**).

**Figure 2.**
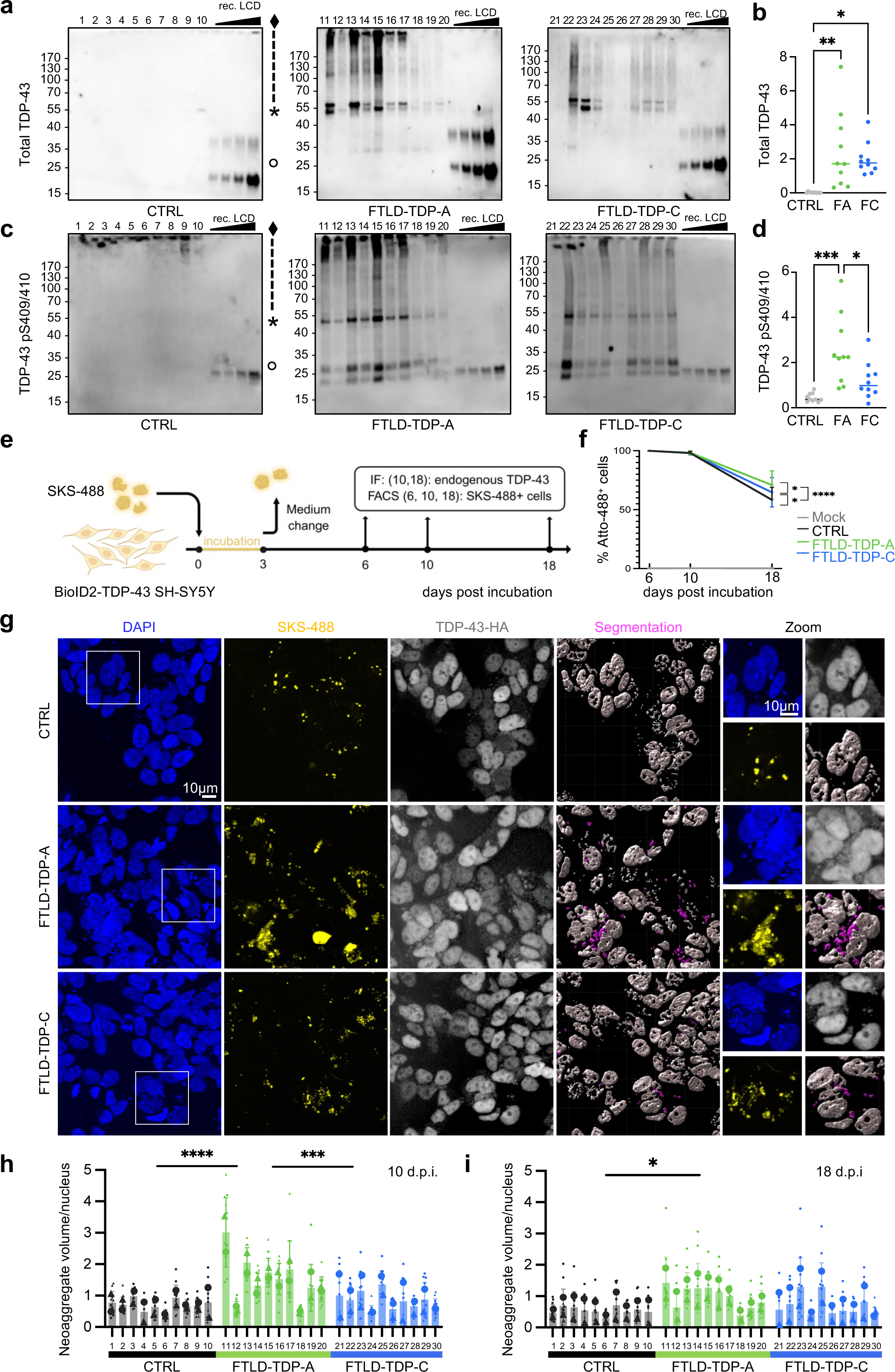
Patient-derived seeds exhibit variable pathological burden and seeding potency. (**a-d**) Biochemical analysis of seeds extracted from postmortem human brain samples with the SarkoSpin method^18^. Seeds were subjected to SDS–PAGE followed by immunoblotting against (**a-b**) total TDP-43 with a cocktail of antibodies against TDP-43 N-terminus and C-terminus and (**c-d**) phosphorylated TDP-43 at position Ser409 or Ser410 (TDP-43 pS409/410) with a commercially available antibody. Note the various TDP-43 species observed in SarkoSpin pellets from FTLD patient brains: hyperphosphorylated/polyubiquitination smear at 70 kDa and above (♦), full length TDP-43 (*) and at the C-terminal fragment of approximately 25 kDa (°). (**b, d**) Quantifications of immunoblots showing the average of two repetitions of the experiment. Statistical analyses (n=10) were performed using a one-way ANOVA comparing the interaction effects of time points and treatment conditions, with Tukey’s multiple comparisons (*p < 0.05, **p < 0.01, ***p < 0.001, ****p < 0.0001). (**e**) Schematic representation of our seeding model. SH-SY5Y cells stably expressing BioID2-TDP-43WT were incubated for 3 days with Atto-488-labeled patient-derived seeds (SKS-488) before removing them by medium exchange. (**f**) Percentage of cells with internalized TDP-43 aggregates measured by FACS after incubation with either seeds from non-neurodegenerative control (CTRL, black), FTLD-TDP-A (green), FTLD-TDP-C (blue) with 5 individuals (n=5) per condition as biological replicates, or without any treatment as a mock control (gray). The lines represent the mean of measurements taken from 3 independent experiments, and error bars represent the standard deviation. Statistical analyses were performed using a two-way ANOVA comparing the interaction effects of time points and treatment conditions, with Tukey’s multiple comparisons (*p < 0.05, ***p < 0.001). (**g**) Representative immunofluorescence images taken at 10 d.p.i. showing nuclei (DAPI, blue), exogenous fluorescently labeled patient-derived aggregates (SKS-488, yellow) and endogenous TDP-43 (TDP-43-HA, white) were use to generate 3D reconstructed objects for segmentation of neoaggregates, which were defined as the colocalized endogenous TDP-43-HA with exogenous aggregates with an overlapping volume of over 30% (segmentation, magenta). Scale bar: 20 μm. (**h, i**) Image quantification displaying the volume of neoaggregates, normalized to the count of cells per field-of-view (FOV) for 10 (**h**) and 18 d.p.i (**i**). Smaller dots represent the value from each FOV, and larger dots represent the mean of measurements taken from two independent experiments, with 10 individuals per condition as biological replicates (n=10). Error bars represent the standard deviation. Statistical analyses were performed for the comparison between CTRL, FTLD-TDP-A and FTLD-TDP-C using a one-way ANOVA with Tukey’s multiple comparisons (*p < 0.05, **p < 0.01, ***p < 0.001, ****p < 0.0001).

To characterize the seeding properties of the patient-derived aggregates, we evaluated their ability to recruit endogenous TDP-43 and form neoaggregates, by quantifying the colocalization of cellular TDP-43-HA with the fluorescently labeled exogenous SarkoSpin seeds. Confocal immunofluorescence images from seeded cells were processed in Imaris to reconstruct three-dimensional objects for quantification (**Fig. 2g, Fig. S3**). At 10 d.p.i. the volume of neoaggregates was significantly higher in cells incubated with FTLD-TDP-A seeds compared to those treated with control or FTLD-TDP-C seeds (**Fig. 2h**), suggesting higher seeding potency of FTLD type A aggregates, as we previously reported^5, 18^. By 18 d.p.i. neoaggregate volume was overall decreased, potentially due to degradation of the exogenous seeds, and/or their Atto-488 label (**Fig. 2i**). High patient-to-patient variability was evident in both FTLD groups across both timepoints (**Fig. 2h, i**). In the BioID2-TDP-43mNLS line, seeding with the same SarkoSpin extracts produced higher rates of neoaggregates in both patient groups and both timepoints, likely driven by the higher cytoplasmic substrate availability (**Fig. S4a-c**). Notably, both the inter-individual variability and the higher seeding potency of the FTLD-TDP-A seeds were similarly evident in both cell lines, validating our results.

### Loss of physiological TDP-43 interactions upon seeding with FTLD-TDP-A and C

After monitoring the internalization rates and seeding efficiencies of all individual brain-derived SarkoSpin extracts (**Fig. 2, Fig. S3-S4**), we proceeded in the identification of changes in TDP-43 interactomes upon seeding to illuminate potentially distinct molecular pathways between the CTRL and patient groups as well as between the two FTLD subtypes. We selected five control individuals and five patient samples for each FTLD-TDP subtype, based on our previous analyses and incubated BioID2-TDP-43WT cells for 10 and 18 d.p.i., prior to isolation of biotinylated proteins and mass spectrometry, as previously done (**Fig. 1**). PCA analysis of the enriched interactomes showed that PC1 primarily separates samples based on the d.p.i., while both PC1 and PC2 allowed sample separation according to their treatment (**Fig. S4d**). When we compared the differential interactome of TDP-43 after seeding with control, FTLD-TDP-A or FTLD-TDP-C seeds at different time points, we identified significantly altered proteins associated exclusively with certain conditions. Notably, at 10 d.p.i., more proteins showed decreased interactions with TDP-43 after seeding with both FTLD-TDP groups compared to controls (**Fig. 3a, b**). This decrease in interaction of TDP-43 with its physiological protein partners suggested the initiation of its loss of function. This was supported by the common FTLD-TDP-A and FTLD-TDP-C biological processes identified via GO enrichment analysis, which pointed to decreased nuclear interactions of TDP-43 with partners in ribonucleoprotein biogenesis and other processes involving binding to nucleic acids (**Fig. 3c, d** and **Fig S5a**). At 18 d.p.i. the alterations of TDP-43 interactome significantly diverged between cells seeded with the two FTLD-TDP subtypes. Specifically, while FTLD-TDP-A-seeded cells, showed extensive interactor disruptions (**Fig. 3e**), FTLD-TDP-C-seeded cells exhibited only minimal significant shifts (**Fig. 3f**), pointing to better recovery and the relative inertness of type C seeds. Interestingly, at this later timepoint, FTLD-TDP-A-seeded cells showed increased interactions with spectrin-binding proteins (**Fig. S5c**), specifically ADD1 (Adducin 1, alpha subunit), which maintains the actin-spectrin cytoskeleton, KIF3A (Kinesin family member 3A), a motor protein important for axonal transport, and EPB41 (Erythrocyte membrane protein band 4.1), which anchors membrane proteins to cytoskeleton in neurons. A connection between TDP-43 skein-like cytoplasmic aggregation and cytoskeletal proteins like actin has been recently highlighted^65^, suggesting that this pathway might be specifically involved in FTLD-TDP-A-seeded aggregation. At both timepoints, only a few differential interactors could be distinguished when comparing FTLD-TDP-A and FTLD-TDP-C (**Fig. 3g, h**), which precluded meaningful GO analyses. However, among these few interactors specifically increased in FTLD-TDP-A-treated cells, we identified proteins associated with neurite growth and cytoskeletal dynamics, including EPB41, CRMP1 (Collapsin Response Mediator Protein 1) and RAB14 (Ras-Related Protein Rab-14), supporting an important role of these processes for seeding outcomes.

**Figure 3.**
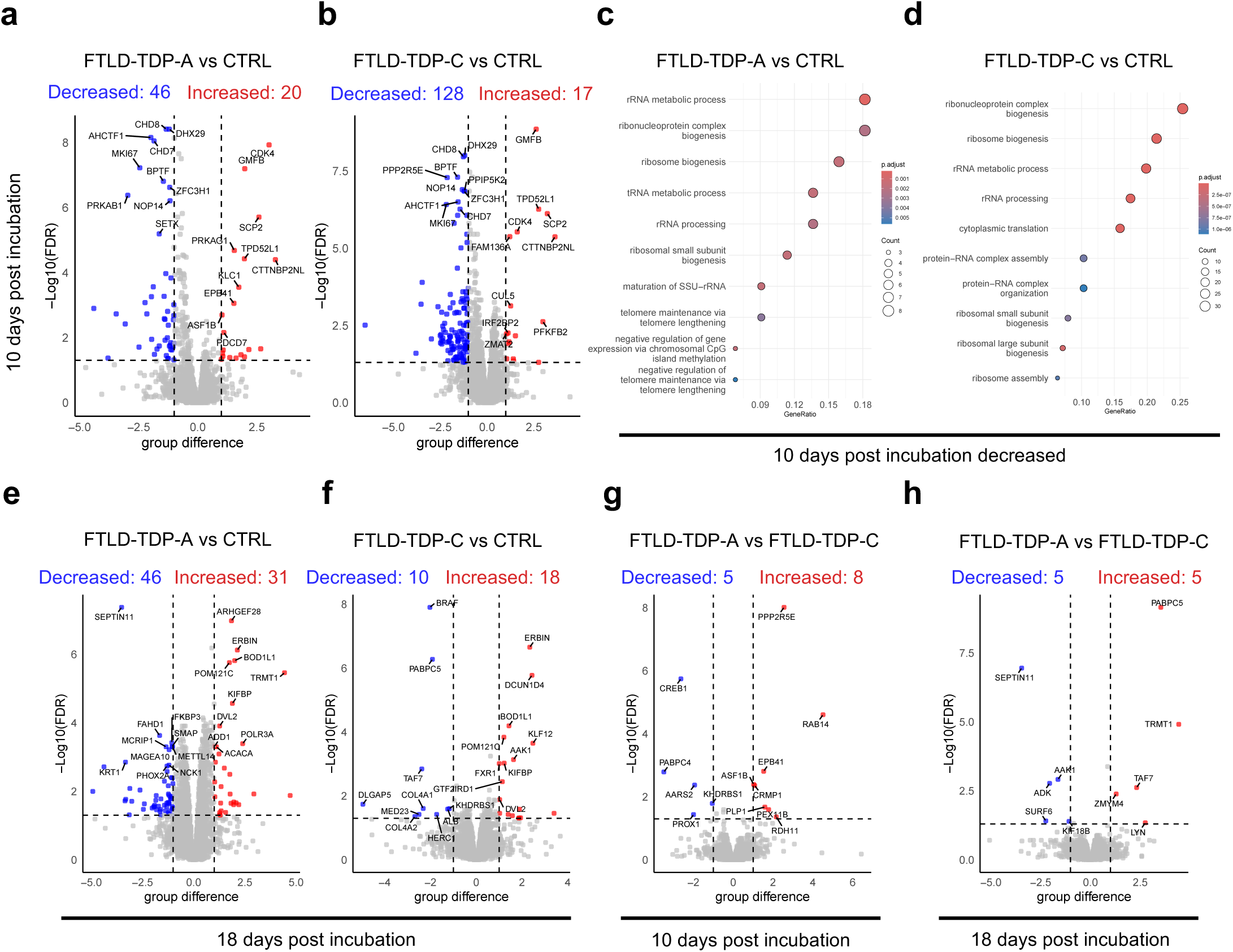
Loss of physiological TDP-43 interactions upon seeding with FTLD-TDP-A and C. Volcano plots show the comparison of experimental conditions at 10 d.p.i for FTLD-TDP-A vs. CTRL (**a**) and FTLD-TDP-C vs. CTRL (**b**) after a differential expression analysis. The dot colors indicate if the interaction of TDP-43 with a protein partner is decreased (blue), increased (red) or not significantly modified (grey) according to the group difference (x-axis) with a threshold of 1-fold and -Log_10_ false discovery rate (FDR) (y-axis) < 0.05. Displayed numbers indicate significantly modified interactions within each comparison. (**c-d**) Enriched Gene Ontology for biological processes showing decreased interactions between FTLD-TDP-A and CTRL (**c**) and FTLD-TDP-C vs. CTRL (**d**) at 10 d.p.i. Dot sizes represent the number of genes in each pathway (count), while dot colors indicate pathway significance based on adjusted p-values (p.adjust). The GeneRatio on the x-axis represents the proportion of DEGs associated with each pathway. (**e-h**) Volcano plots analyzed as described in a-b showing comparisons of FTLD-TDP-A vs. CTRL (**e**) and FTLD-TDP-C vs. CTRL (**f**) at 18 d.p.i., and FTLD-TDP-A vs. FTLD-TDP-C at 10 d.p.i (**g**), or 18 d.p.i. (**h**).

### Seeded aggregation with patient-derived extracts induces TDP-43 loss of function

In various neurodegenerative diseases, TDP-43 loss of function leads to devastating consequences^11, 12^ in affected neurons through widespread splicing dysregulation^17^ and aberrant cryptic exon incorporation^4, 7, 14, 15^. Previous work demonstrated a mechanistic link between cytoplasmic TDP-43 aggregation induced by external TDP-43 fibrils and its loss of function^43, 44^. To track and quantify TDP-43 loss of function in living cells, we employed a TDP-43 splicing regulation reporter (TDP-REG)^47^, which fluorescently labels cells lacking functional TDP-43 in red. SH-SY5Y cells stably expressing TDP-REG^43, 44^ were incubated with Atto-488-labeled patient-derived seeds and analyzed by immunofluorescence and high-content confocal imaging at 10 and 18 d.p.i. (**Fig. 4a** and **Fig. S6**). The majority of mScarlet-positive cells contained large inclusions with recruited endogenous TDP-43, consistent with our previous study using recombinant TDP-43 fibrils^44^. Notably, large inclusions with filamentous or compact morphology were evident in seeded cells, containing both the exogenous, patient-derived material and recruited endogenous TDP-43 (**Fig. 4a** and **Fig. S6**), resembling the morphologies previously reported in patient brains^38^.

**Figure 4.**
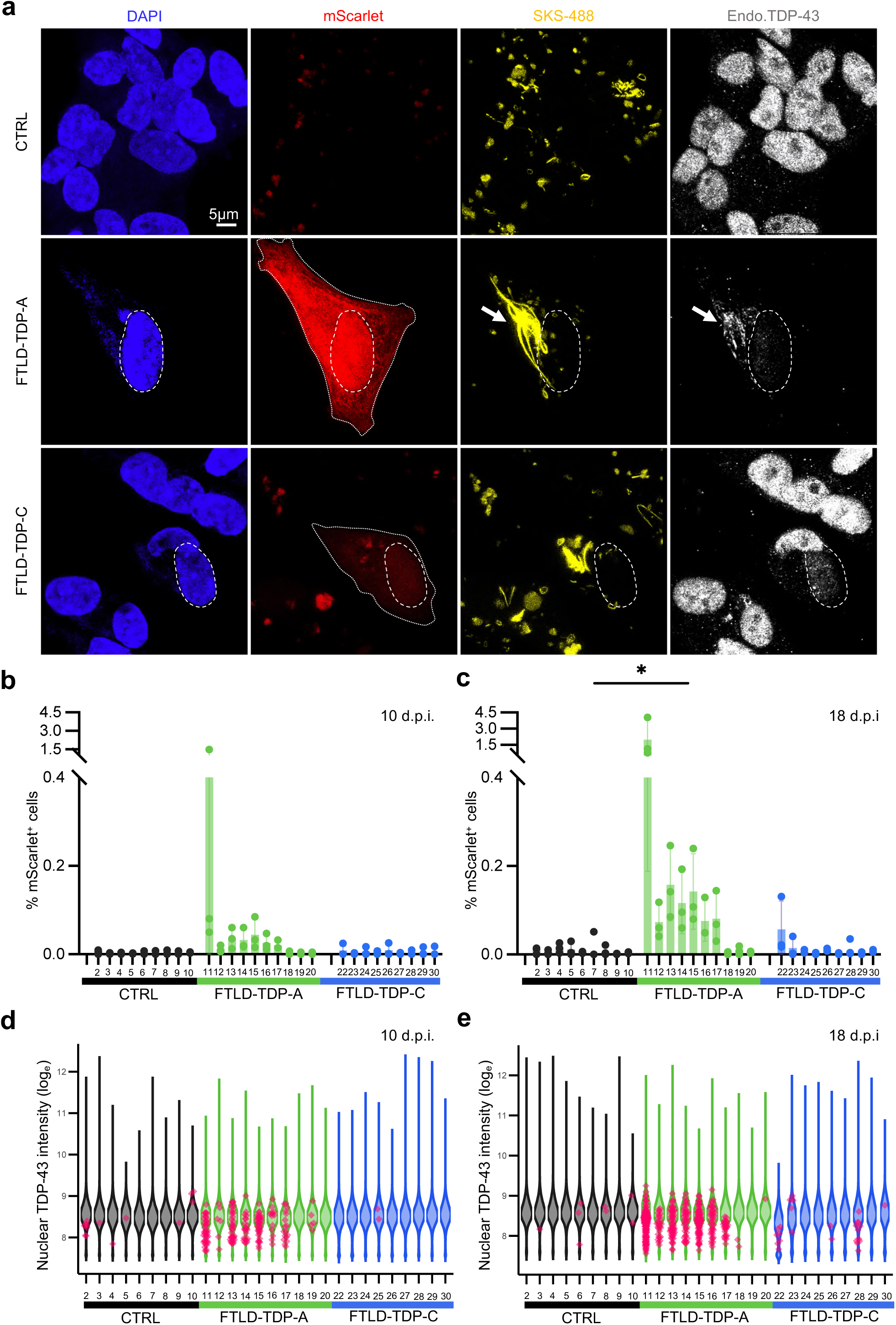
Seeded aggregation with patient-derived extracts induces TDP-43 loss of function. (**a**) Representative immunofluorescence images taken at 18 d.p.i. Dotted lines delineate the outline of mScarlet-positive cells, whereas dashed lines mark the nuclear region of the same cell that displayed reduced endogenous TDP-43 intensity. The white arrowhead marks an example of internalized filamentous TDP-43 extracted from a FTLD-TDP-A brain with the recruitment of endogenous TDP-43 within an mScarlet-positive cell. Scale bar: 5 μm. (**b, c**) Image quantification displaying the percentage of mScarlet-positive cells out of total cells analyzed on 10 and 18 d.p.i, respectively. Data acquired from three independent experiments, with 10 individuals per group as biological replicates (n=10). Error bars represent the standard deviation. Statistical analyses were performed for the comparison between CTRL, FTLD-TDP-A and FTLD-TDP-C using a one-way ANOVA with Tukey’s multiple comparisons (*p < 0.05). (**d, e**) Representative plots of log-transformed integrated TDP-43 nuclear intensity at 10 (**d**) and 18 d.p.i. (**e**). The violin shapes denote the distribution of TDP-43 nuclear intensity of mScarlet-negative cells from a representative replicate with juxtaposed red dots, representing TDP-43 nuclear integrated intensity of mScarlet-positive cells identified and quantified in each condition.

Image-based quantification at 10 d.p.i. showed a small percentage of mScarlet-positive cells, in some samples of the FTLD-TDP-A-seeded, but not the FTLD-TDP-C-seeded, group, yet this did not reach statistical significance (**Fig. 4b**). In contrast, at 18 d.p.i., the percentage of mScarlet-positive cells increased in most samples of the FTLD-TDP-A-seeded group and few of the FTLD-TDP-C, with the former reaching statistical significance compared to the CTRL group (**Fig. 4c**). The distribution of nuclear TDP-43 intensity showed that mScarlet-positive cells generally had reduced nuclear TDP-43 compared to the overall population (**Fig. 4d, e**), with this reduction being most evident at 18 d.p.i., consistent with the higher proportion of mScarlet-positive cells at this time point (**Fig. 4d, e**). This increase in mScarlet-positive cells over time (**Fig. 4b-e**), contrasted with the observed decrease in exogenous aggregates (**Fig. 2h, i**) and indicates a propagating cascade, eventually leading to loss of TDP-43 function.

### Inferential analysis reveals the TMEM106B fibrillar core as a TDP-43 pro-seeding factor

The high inter-patient variability in neoaggregate formation (**Fig. 2**) and induced loss of function (**Fig. 4**), prompted us to interrogate the determinants of these seeding outcomes. Various TDP-43 species in patient-derived extracts, including phosphorylated TDP-43 and C-terminal fragments have been proposed to be seeding-competent^5, 19, 20, 43, 44^. Our biochemical characterization of SarkoSpin pellets showed enrichment of total TDP-43 and pTDP-43 in FTLD-TDP-A and FTLD-TDP-C samples, but with a great level of heterogeneity among individual patients (**Fig. 2a-d**). Control samples were mostly devoid of total TDP-43, confirming that the SarkoSpin method efficiently removes the spurious physiological protein while enriching for the pathological species (**Fig 2a-d**). We further classified the detected TDP-43 protein bands by their characteristic signatures into high-molecular weight species with polyubiquitination or hyperphosphorylation (70+kDa), full-length (FL), and C-terminal fragments (CTF) (**Fig. 2a, c**). We considered two additional factors that were recently shown to contribute to the neuropathological and clinical heterogeneity of these FTLD-TDP subtypes. First, ANXA11 was recently found to be co-assembled with TDP-43, forming distinctive heterotypic amyloid filaments specifically in FTLD-TDP-C patients^28^ characterized by distinct clinical presentation and slow disease progression^18, 26^. Indeed, independent neuropathological analyses reported co-deposition of pathological TDP-43 with ANXA11, specifically in FTLD-TDP-C patients^29–31^. Our group previously identified ANXA11 among the insoluble proteins in the SarkoSpin pellets of FTLD-TDP-C patients, but not other subtypes, with an unbiased proteomic approach^18^. Indeed, our biochemical characterization of SarkoSpin pellets of the current cohort, confirmed the presence of ANXA11 fragment, in FTLD-TDP-C patient extracts, but not in FTLD-TDP-A or controls (**Fig. 5a, b**). Second, the transmembrane lysosomal protein TMEM106B was shown to assemble into amyloid filaments in aging brains and in multiple neurodegenerative conditions^32–34^. While the role of these filaments in disease remains unclear, SNPs in the gene encoding TMEM106B were linked to the risk of developing FTLD, with certain protective alleles linked to extended survival after onset^35–37^. This genetic link is particularly strong in GRN and C9orf72 mutation carriers^66^ who most frequently present with FTLD-TDP-A neuropathology^38^. Using an antibody specific to the TMEM106B fibrillar core, we detected variable levels of the corresponding fibrils in the insoluble fraction from all samples (**Fig. 5c, d**) as previously reported^32–34^, with some FTLD-TDP-A samples (FA-11, FA-15-17) showing markedly higher levels.

**Figure 5.**
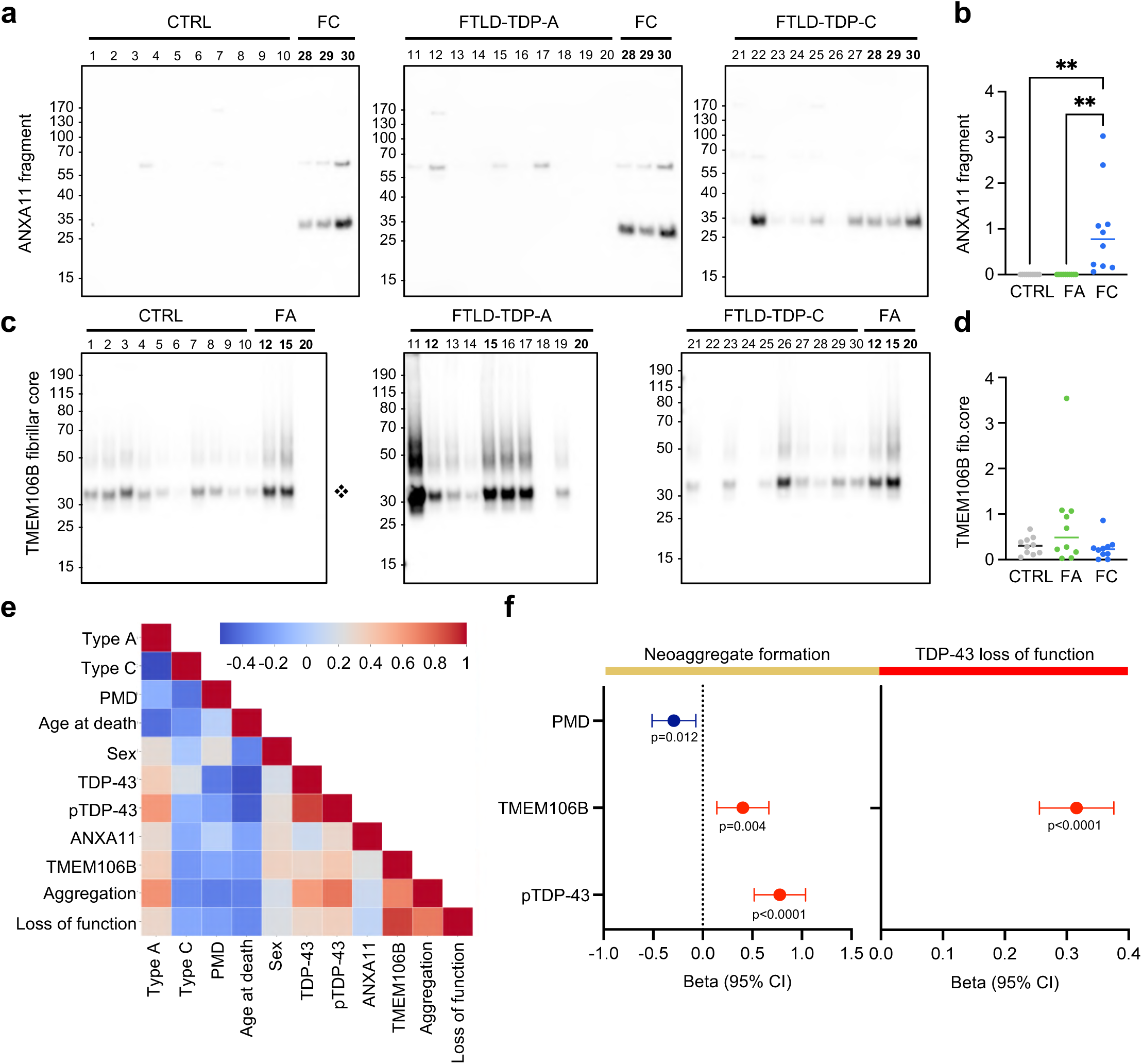
Inferential analysis reveals the TMEM106B fibrillar core as a TDP-43 pro-seeding factor. (**a-d**) Biochemical analysis of seeds extracted from postmortem human brain samples with the SarkoSpin method^63^. Seeds were subjected to SDS–PAGE followed by immunoblotting against (**a**) total and fragmented ANXA11 observed at ∼35kDa and (**c**) TMEM106B fibrillar core observed at ∼ 30kDa. (**b, d**) Quantifications of immunoblots. For TMEM106B, the data displays the average of two repetitions of the experiment. Statistical analyses (N=10) were performed using a one-way ANOVA comparing the interaction effects of time points and treatment conditions, with Tukey’s multiple comparisons (*p < 0.05, **p < 0.01, ***p < 0.001, ****p < 0.0001,). (**e**) Pairwise correlations of clinical and biochemical variables of the cohort of samples used for seeding, along with their capacity of inducing neoaggregates and loss of TDP-43 function, were assessed using Pearson’s correlation coefficient. Statistical significance of correlations was evaluated using two-tailed t-tests. Results were visualized as a correlation heatmap with the color scale representing correlation strength and direction. (**f**) Final multiple linear regression models linking quantified patient variables to TDP-43 neoaggregate formation and loss of function. The figure presents the final models obtained through backward variable selection, illustrating how quantified variables from individual patients contribute to neoaggregation formation (left graph) or the induction of TDP-43 loss of function (right graph). Regression coefficients (β) are shown with corresponding 95% confidence intervals (CI). The models account for a large proportion of variance, with adjusted R² values of 0.788 for neoaggregation and 0.801 for TDP-43 loss of function.

Using all available biochemical variables (**Fig. 2a-d** and **Fig. 5a-d**) and clinical data for these patients (**Table S3**), we applied multiple linear regression to determine whether any variables showed a linear association with the seeding outcomes. As outcomes, we considered neoaggregate formation at 10 d.p.i and TDP-43 loss of function at 18 d.p.i, corresponding to the peak of each effect. Since the relative seeding capacity of individual patient materials remained consistent across time points, we selected a single representative time point as the primary outcome to avoid redundancy, while incorporating both clinical factors and biochemical variables in the analysis. We initiated a full model containing all quantified variables, with backward stepwise selection to identify the best fitting model with non-redundant variables^67^. In the final model, the load of pTDP-43 and TMEM106B fibrillar core showed a significant positive contribution to neoaggregate formation (**Fig. 5e, f** and **Fig. S7**). Conversely, the postmortem delay (PMD), showed a significant negative association, possibly due to increased degradation of the pathological seeds during sample preparation. Altogether, the model explains 78% of the variability in the response. Surprisingly, the same analysis using TDP-43 loss of function as the outcome, identified TMEM106B fibrillar core as the only significant variable in the final model, independently accounting for 80% of the variability in the data (**Fig. 5e, f** and **Fig. S7**). Overall, our data indicate that the presence of the TMEM106B fibrillar core promotes loss of TDP-43 function, after seeding with patient-derived aggregates.

### Lysosomal escape unlocks TDP-43 seeding in human neurons

How could TMEM106B fibrils contribute to the induction of TDP-43 loss of function? To answer this question, and validate our observations in human neurons, we installed the previously characterized TDP-REG reporter^44, 47^ in human induced pluripotent stem cells (hiPSCs) harboring the Dox-inducible *NGN2* cassette for their controlled differentiation into cortical excitatory neurons^68^. Upon lentivirus-mediated short-hairpin *TARDBP* knockdown (KD), almost 100% of neurons turned red due to TDP-REG activation, confirming successful line generation and the expected reporter activation (**Fig. 6a-c**). Notably, untreated neurons showed no mScarlet signal, validating the tightness of the system.

**Figure 6.**
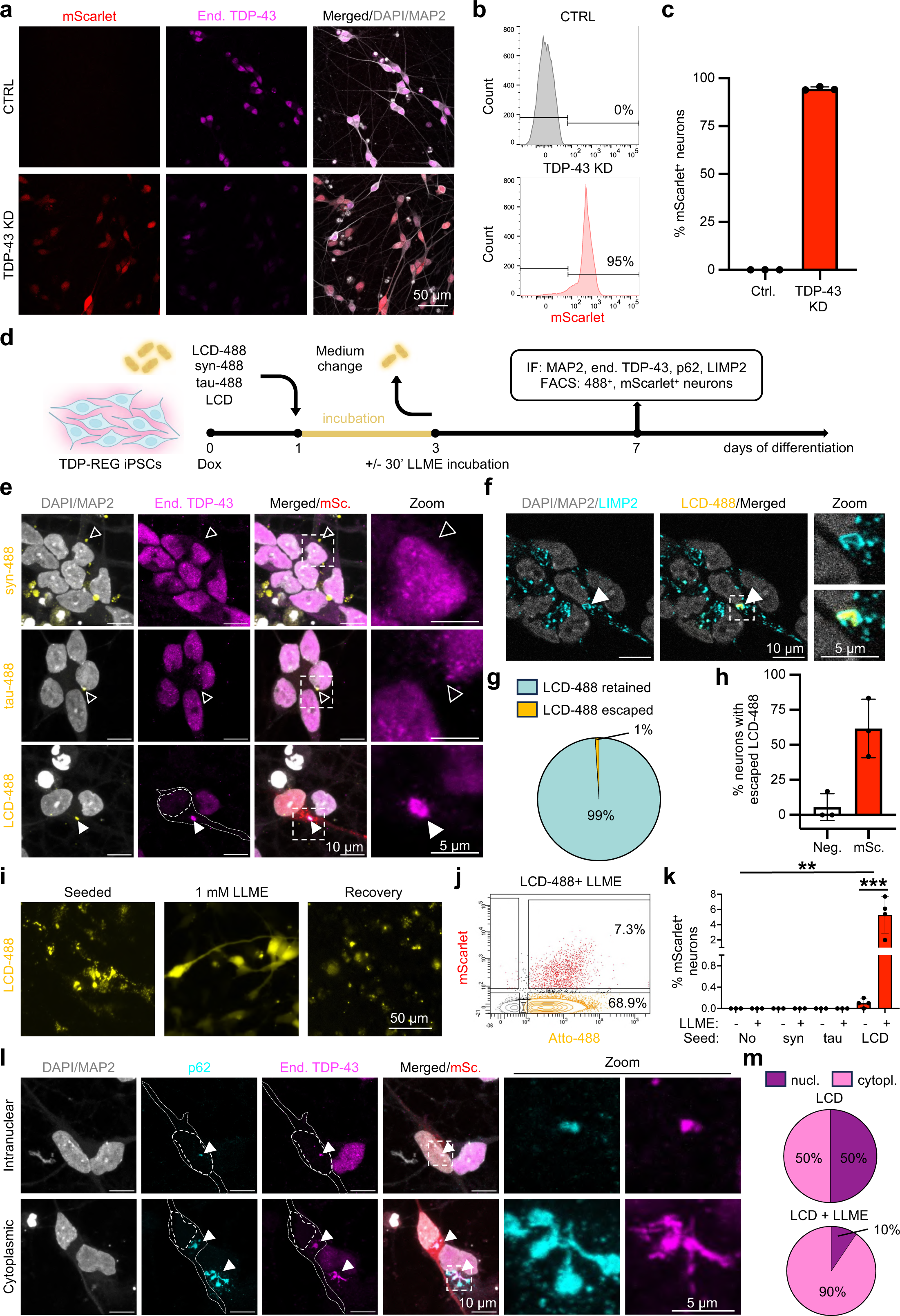
Lysosomal escape unlocks TDP-43 seeding in human neurons. (**a-c**) Immediately after plating, TDP-REG *NGN2* iPSCs were transduced with a lentivirus carrying shRNA against *TARDBP*. Neurons were differentiated for 7 days before (**a**) immunostaining for TDP-43 (magenta; scale bar: 50 µm), or (**b**) fluorescent activated cell sorting (FACS) for mScarlet-positive neurons, which are detectable only after TDP-43 knockdown (KD). (**c**) FACS quantification of three independent experiments. Each single dot represents the percentage of mScarlet-positive neurons for each experiment (n=3). Error bars show standard deviation (SD). (**d**) Schematic representation of our seeding model using TDP-REG *NGN2* iPSCs. (**e**) Representative immunofluorescence images (day 7) revealed mScarlet-positive neurons (red, third column) after treatment with LCD-488, but not a-synuclein or tau. mScarlet-positive neurons showed cytoplasmic inclusions composed of an LCD-488-positive core (yellow) colocalizing with endogenous TDP-43 recognized by an antibody against the N-terminal domain of TDP-43 (magenta), shown also in the zoomed insets (dotted-line squares). Notably, TDP-43 cytoplasmic aggregation is accompanied by the reduction of its nuclear levels. Dotted lines show the body of the mScarlet-positive neuron and the nuclear area overlayed on the TDP-43 signal. White arrowheads mark exogenous LCD-488 aggregates co-localizing with endogenous TDP-43, while empty arrowheads mark a-syn-488 and tau-488 aggregates that do not recruit endogenous TDP-43. Scale bar: 10 µm and 5 µm in the zoomed images (n=3). (**f**) Representative immunofluorescence image (day 7) showing exogenous LCD-488 fibrils (yellow) surrounded by LIMP2 signal (cyan). Scale bar: 10 µm and 5 µm in the zoomed images (n=3). (**g**) Pie chart showing the quantification of LCD-488 fibrils retained in or escaped from the lysosomes. (**h**) Graph showing the percentage of mScarlet-positive or -negative neurons containing escaped LCD-488 fibrils (n=3; manual quantification for both **g** and **h;** 10 FOVs analyzed per replicate containing at least one mScarlet-positive neuron). Each single dot represents the mean percentage of neurons with escaped aggregates for each of the three independent experiments. Error bars show SD. (**i**) Live imaging of NPCs seeded with 200 nM LCD-488 fibrils showing the reversible effect of LLME (30’ incubation with 1 mM) on fibril compartmentalization. Images were acquired before LLME (left panel), immediately after incubation (central panel), and 24 hours after LLME removal (right panel). Scale bar: 50 µm. (**j**) FACS contour plot of TDP-REG *NGN2* neurons (day 7) showing Atto-488-positive and mScarlet-positive neuron distribution after seeding with 200 nM LCD-488 combined with LLME treatment. (**k**) Percentages of mScarlet-positive neurons upon seeding with different fibrils with or without LLME treatment (n=4 for LCD seeds, n=3 for the rest). Each dot represents the percentage of mScarlet-positive neurons in one independent experiment. Error bars show SD. Two-way ANOVA with Šidák multiple-comparisons test (**p < 0.01, ***p < 0.001). (**l**) Representative immunofluorescence images (day 7) showing mScarlet-positive neurons seeded with 200 nM of unlabeled LCD displaying intranuclear (top panels), or cytoplasmic (lower panels) inclusions positive for endogenous TDP-43 (magenta) and p62 (cyan). Dotted lines show the body and nucleus of the mScarlet-positive neurons. Arrowheads mark p62 signal co-localizing with endogenous TDP-43, also shown in the zoomed insets (dotted-line squares). Scale bar: 10 µm and 5 µm in the zoomed images. (**m**) Pie charts show the quantification of nuclear or cytoplasmic inclusions in neurons seeded with LCD fibrils with or without LLME treatment (n=3; 10 FOVs analyzed per replicate containing at least one mScarlet-positive neuron).

To assess the tractability and specificity of seeding outcomes in these neurons, we incubated them with *in vitro* obtained fibrils, composed of the TDP-43 low-complexity domain (LCD), which, we previously showed to induce cytoplasmic aggregation of endogenous TDP-43 and its consequent loss of function in neuron-like cells^44^. We incubated neuronal precursor cells (NPCs, differentiation day 1) for two days with 200 nM unlabeled or Atto-488 labeled TDP-43 fibrils (LCD-488), and performed FACS and immunofluorescence analyses at day 7, when cells were differentiated into neurons. As controls we incubated NPCs with the same concentration of fluorescently labeled α-synuclein and tau K18 fibrils^69^ (**Fig. 6d**). TDP-43 recruitment and consequent loss of function occurred exclusively in neurons exposed to LCD fibrils but not α-synuclein or tau K18 species (**Fig 6e, Fig. S8a, b**). Notably, the nuclei of mScarlet-positive neurons showed depletion of endogenous TDP-43 – specifically recognized by an antibody directed against the N-terminal domain that is excluded in the LCD-488 fibrils – and colocalization with the exogenous LCD-488 fibrils in the cytoplasm (**Fig. 6e**).

In agreement with what we previously reported in neuron-like cells^44^, most neurons (∼90%) internalized LCD-488 fibrils, yet only a small population (∼0.1%) developed loss of TDP-43 function and activated the TDP-REG to generate mScarlet-positive neurons (**Fig. S8b**). Since in our seeding model fibrils are freely internalized by cells, the observed low recruitment efficiency of internalized TDP-43 fibrils may reflect lysosome sequestration with limited access to the cytoplasmic compartment. To test this hypothesis, we stained seeded neurons with an antibody recognizing the lysosomal membrane protein LIMP2, which revealed lysosomal sequestration of almost all internalized LCD-488 fibrils (**Fig. 6f, g**). Notably, lysosome-escaped (i.e. not LIMP2-surrounded) LCD-488 fibrils were mainly contained in mScarlet-positive neurons (**Fig. 6h**), suggesting that fibril escape is needed to induce efficient TDP-43 recruitment and consequent loss of function. Therefore, we wondered if a brief injury of the lysosomal membrane might enhance seeding outcomes. To test this, we used the lysosomotropic agent L-leucyl-L-leucine, methyl ester (LLME), which was shown to enter lysosomes through receptor-mediated endocytosis and to be converted by dipeptidyl peptidase I to a hydrophobic polymer with membranolytic activity, resulting in the transient permeabilization of lysosomal membranes^70^. After 2 days of incubation with 200 nM TDP-43 LCD fibrils, cultures were treated with 1 mM LLME for 30 minutes to transiently permeabilize lysosomal membranes after fibril internalization. We selected 1 mM LLME concentration based on prior evidence of efficacy and minimal toxicity in cell lines^70^ and neurons^71^. Live-cell imaging confirmed the activity of the LLME treatment, as it showed the redistribution of internalized LCD-488 fibrils from a dot-like pattern, consistent with lysosomal compartmentalization prior to treatment, to a diffused signal covering the whole cell within 30 minutes after treatment, an effect that was reversed 24 hours later (**Fig. 6i**). Excitingly, LLME-mediated lysosome permeabilization dramatically increased the number of mScarlet-positive neurons by up to ∼70 times (**Fig. 6j**), without affecting seeding specificity, since LCD but not α-synuclein and tau K18 fibrils induced TDP-43 loss of function (**Fig. 6k**). Moreover, another predominantly nuclear RNA-binding protein, FUS was unaffected in mScarlet-positive neurons (**Fig. S8c, d**), confirming the specificity of this induced cytoplasmic localization of TDP-43. An extensive image-based characterization of TDP-43 pathology, which we defined as aggregates composed of recruited endogenous TDP-43 co-labeled by p62, revealed two different phenotypes. A first set of affected neurons showed cytoplasmic TDP-43 aggregates, while a second one contained intranuclear inclusions (**Fig. 6l, m** and **Fig. S8e, f**). The latter are exclusive to neurons and were not observed in seeded SH-SY5Y neuron-like cells^44^. Importantly, the occurrence of intranuclear TDP-43-positive inclusions is extensively described in ALS and FTLD pathological reports^1, 2, 72^, along with the description of intranuclear p62 deposits, which variably co-localize with TDP-43^73, 74^. Interestingly, lysosome permeabilization affected the relative frequency of these phenotypes, increasing the number of cytoplasmic aggregates from 50% to 90% (**Fig. 6m**), via an unknown mechanism.

### TMEM106B promotes TDP-43 seeding with patient-derived TDP-43 fibrils

Our data revealed that the presence of the TMEM106B fibrillar core within the seeds (**Fig. 5c, d**) or a transient lysosomal membrane rupture post exposure to the seeds (**Fig. 6**), enhanced the seeding outcomes in human neuron-like cells and in neurons, respectively. To understand if these observations are linked and to define the contribution of TMEM106B and lysosomal escape in TDP-43 seeding efficiency, we seeded TDP-REG *NGN2* human neurons with SarkoSpin extracts of the same cohort of 10 CTRL, 10 FTLD-TDP-A and 10 FTLD-TDP-C patient samples and quantified mScarlet-positive neurons. As we did with *in vitro*-generated fibrils (**Fig. 6d**), we exposed neurons to SarkoSpins with or without incubation with LLME (**Fig. 7a**), prior to FACS analysis. To verify that the measured mScarlet signal was not affected by the intrinsic autofluorescence of brain-derived material, we applied the same treatment to non-TDP-REG *NGN2* human neurons using as seeds our best FTLD-TDP-A seeder (FA 11) and one CTRL, confirming the specificity of the mScarlet signal to the activation of the reporter and the clear distinction with the detectable autofluorescence of the SarkoSpin extracts (**Fig. S9a-c**). While we only detected seeding activity in neurons treated with FTLD-TDP-A, but not FTLD-TDP-C or CTRL samples (**Fig. 7b**), not all FTLD-TDP-A samples showed the same seeding efficiency. In fact, the seeding outcome of individual extracts could be ranked according to their respective TMEM106B load (**Fig. 7b**), as biochemically quantified (**Fig. 5c, d**). In line with our observations with recombinant LCD fibrils, the combination with LLME incubation enhanced the number of neurons with seed-induced TDP-43 loss of function (**Fig. 7c**). Importantly, also in this case the combination of seeding with LLME increased the occurrence of cytoplasmic TDP-43 deposits (**Fig. 7d, e, Fig. S9d**). Image-based quantification of mScarlet-positive seeded neurons showed incomplete or absent LIMP2 signal surrounding those SarkoSpin aggregates that efficiently recruited endogenous TDP-43 (recruiters) compared to inactive ones that were fully sequestered within lysosomal membranes (**Fig 7f, g, Fig. S9e**). Interestingly, FTLD-TDP-A seeders in the absence of LLME-mediated lysosomal permeabilization showed higher percentages of mScarlet-positive neurons (ranging from ∼2 to 15%) (**Fig. 7b**) compared to neurons exposed to recombinant LCD fibrils (0.1%) (**Fig. 6k, Fig. S8b**), indicating that patient-derived seeds possess different properties and are more potent than pure *in vitro*-obtained LCD fibrils. The observed correlation between seeding efficiency and TMEM106B fibril burden suggests that one potentiator of the enhanced seeding outcomes could be the presence of TMEM106B fibrils. Overall, our work identifies a novel role of lysosomal injury and TMEM106B fibrils in TDP-43 aggregation and induced loss of function.

**Figure 7.**
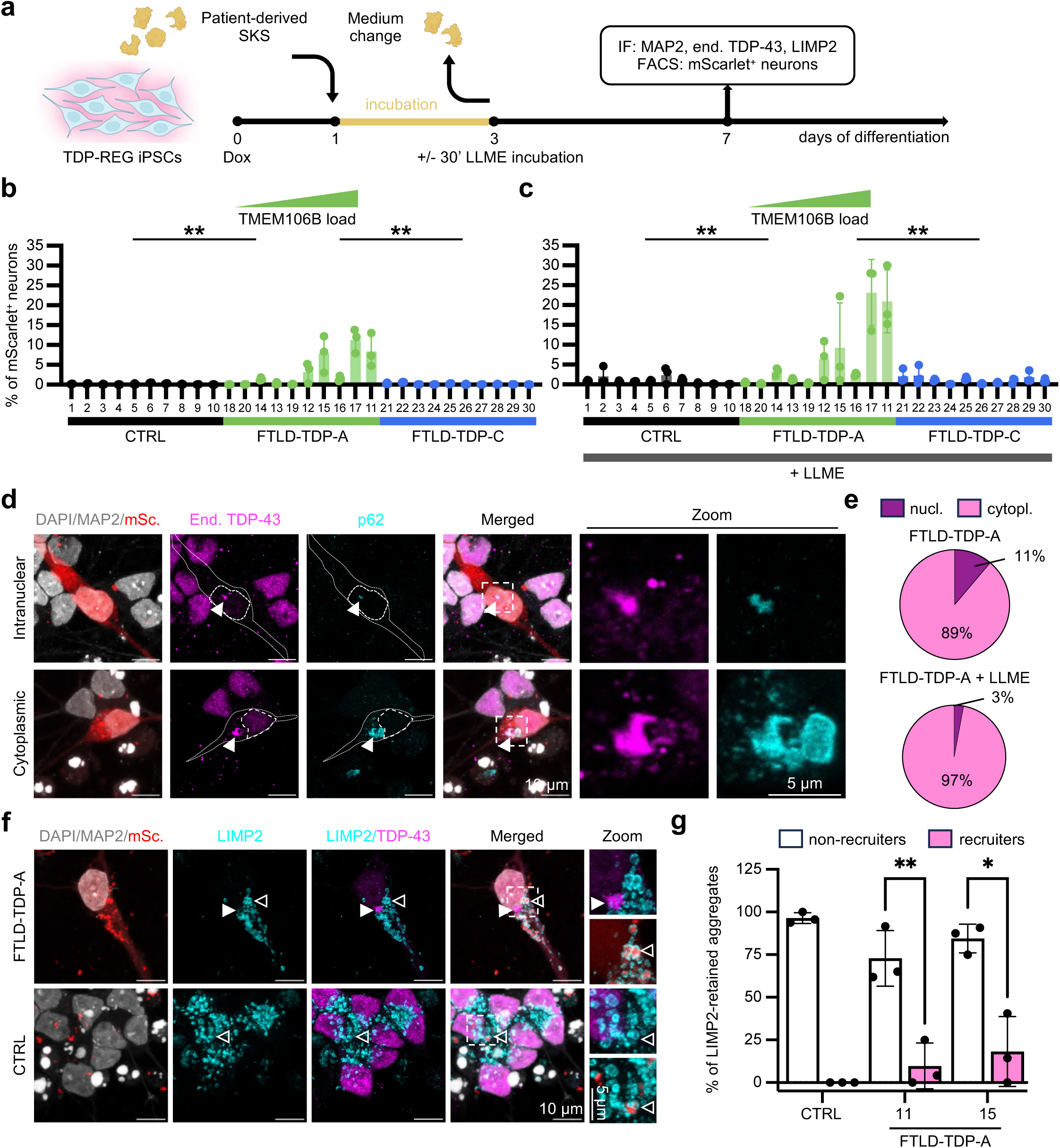
TMEM106B promotes TDP-43 seeding with patient-derived TDP-43 fibrils. (**a**) Schematic representation of our seeding model in TDP-REG *NGN2* neurons. (**b-c**) Percentages of mScarlet-positive neurons after incubation with SKS seeds from different patients without (**b**) or with (**c**) LLME treatment. Each single dot represents the percentage of mScarlet-positive neurons for each of the three independent experiments (n=3). Error bars show standard deviation (SD). Statistical analyses comparing groups were performed using two-way ANOVA with Tukey’s multiple comparisons **p < 0.01). FTLD-TDP-A samples are ordered according to increasing TMEM106B content as quantified by western blot in Fig. 5c, d. (**d**) Representative immunofluorescence images (day 7) showing mScarlet-positive neurons after seeding with FTLD-TDP-A SKS material (nr.11) displaying intranuclear (top panels) and cytoplasmic (lower panels) inclusions positive for endogenous TDP-43 (magenta) and p62 (cyan). Dotted lines show the body and nucleus of the mScarlet-positive neurons. Arrowheads mark p62 signal co-localizing with endogenous TDP-43, also shown in the zoomed insets (dotted-line squares). Scale bar: 10 µm and 5 µm in the zoomed images. (**e**) Pie charts show the quantification of nuclear or cytoplasmic inclusions in neurons seeded with FTLD-TDP-A SKS with or without LLME treatment (n=3; 10 FOVs analyzed per replicate containing at least one mScarlet-positive neuron. Phenotypes were assessed in two FTLD-TDP-A: nr. 11 and 13; pie charts show the average of the two samples). (**f**) Representative immunofluorescence images (day 7) showing an mScarlet-positive neuron after seeding with FTLD-TDP-A SKS material (nr.11) or neurons incubated with CTRL SKS (nr.2), both with LLME treatment (lower panel). The mScarlet-positive neuron shows reduced endogenous nuclear TDP-43 (magenta) and cytoplasmic aggregation. LIMP2 staining (cyan) shows lysosomal sequestration of exogenous SKS material (red, autofluorescent in the 568 channel used for mScarlet). White arrowheads mark endogenous TDP-43 not surrounded by LIMP2 signal, while empty arrowheads mark SKS material sequestered within lysosomal compartment. Scale bar: 10 µm and 5 µm in the zoomed images (n=3). (**g**) Quantification of autofluorescent patient-derived SKS material retained within the lysosomal compartment (n=3; 10 FOVs analyzed per replicate containing at least one mScarlet-positive neuron). All mScarlet-positive cells per FOV were examined for the presence of SKS material and scored for colocalization with endogenous TDP-43 (recruiters) and LIMP2 (non-recruiters) by manual slice-by-slice inspection of the z-stacks. In the CTRL patient sample, only mScarlet-negative cells were analyzed, and only non-recruiting SKS aggregates were present and quantified for their colocalization with LIMP2. See also Fig. S9e. Each dot represents the mean percentage of LIMP2-retained aggregates for each of the three independent experiments. Error bars show SD. Statistical analyses comparing groups were performed using two-way ANOVA with multiple comparisons (*p < 0.05, **p < 0.01).

## Discussion

Here, we employed our quantitative seeding model^44^ to explore the determinants of patient-derived TDP-43 aggregate internalization, neoaggregate formation, and subsequent TDP-43 loss of function and to identify alterations in the interactome of endogenous TDP-43 during this process. Leveraging a cohort of 30 postmortem brain samples, we assessed if FTLD-TDP types A and C involve distinct molecular pathways. To our knowledge, this represents the largest seeding study with human brain extracts from patients with TDP-43 proteinopathies to date. By integrating this approach with proximity proteomics, we uncovered TDP-43 interactors that varied with seeding conditions over time. Decreased interactions with proteins that function in ribonucleoprotein complex biogenesis were common to both FTLD-TDP-A and FTLD-TDP-C compared to controls early post treatment (10 d.p.i.), consistent with loss of the physiological TDP-43 interactors. At the later time point (18 d.p.i.), these reduced physiological interactions persisted in FTLD-TDP-A, which also showed increased loss of function based on TDP-REG quantifications, indicating that disruption of these pathways intensifies as pathology progresses. In contrast, FTLD-TDP-C-treated cells recovered most physiological interactions over time and showed minimal induction of neoaggregation and loss of function at the same timepoint, in line with the previously reported lower seeding potency^5, 19, 20^ and neurotoxicity^18–20^ of this disease subtype.

Beyond revealing reduced interactions with canonical TDP-43 partners, our unbiased proteomics identified increased binding of TDP-43 to actin-associated proteins, upon treatment with patient brain extracts initially from both FTLD subtypes, but persisting only in the FTLD-TDP-A potent seeders. This finding aligns with a recent report showing TDP-43-actin co-aggregation in cell models and in patient samples, especially when TDP-43 aggregated display a characteristically filamentous or skein-like morphology^65^. In line with this, we identified several seeding-competent filamentous aggregates within FTLD-TDP-A-treated cells, albeit most aggregates appeared granular and compact. Interestingly, the interaction of TDP-43 with actin-associated proteins and the development of filamentous aggregates^65^ was reported to be associated with reduced NTD-mediated TDP-43 oligomerization^75–77^, suggesting that seeding with patient-derived extracts in our system may induce TDP-43 monomerization prior to aggregation. Notably, in FTLD-TDP-A-treated cells, interactions between endogenous TDP-43 and several vesicle-transport proteins (VPS26C, SEC24D, KIF3A, KIFBP, ARL10, EPB41, TMEM160) increased at 18 d.p.i., the time point when loss of function peaked. This association suggests that disruptions in vesicle-mediated transport contribute to the pathological cascade leading to aggregation-linked loss of function.

While we specifically enriched pathological TDP-43 species with SarkoSpin prior to treatment, patient-derived extracts remain highly heterogeneous, both in the relative amount of TDP-43 proteoforms, and in the overall insoluble proteome, as we previously reported^18^. We leveraged this heterogeneity and performed statistical modeling to dissect key factors that determine the seeding outcomes. Our analysis revealed that the most significant determinant of neoaggregate formation, is the presence of pathological TDP-43, supporting a dose effect of seeding-competent species, which may be partially degraded in samples with longer postmortem delay, as the latter showed a significant negative correlation in our analysis. Surprisingly, a critical factor enhancing neoaggregate formation was the presence of TMEM106B fibrillar core, which emerged as the sole determinant of the seed-induced loss of function, an observation that we experimentally validated via seeding with the same cohort of 30 postmortem samples in human neurons.

While we do not know the mechanism of this TMEM106B-mediated boost of seeding outcomes, we hypothesized that it is linked to lysosomal function or disruption thereof, since TMEM106B is a type II transmembrane lysosomal protein, previously shown to regulate lysosomal trafficking in neurons^78^ and to facilitate efficient lysosomal organization and protein degradation^79^. Moreover, TMEM106B has been implicated as an important contributor to FTLD pathogenesis from both genetic^35–37, 80, 81^ and neuropathological evidence^32–34^, though the link between the two phenomena remains unclear. Importantly, mutations in the *GRN gene* encoding for the lysosomal protein progranulin^82^ are causative of FTLD-TDP^83–85^, with hypothesized synergisms with TMEM106B pathology^86^. Several ALS- and FTLD-causing genetic mutations cluster within the endo-lysosomal and autophagy pathways, including *VCP*^87, 88^, *CHMP2B*^89^, *SQSTM1/p62*^90, 91^ and the *C9ORF72* hexanucleotide repeat expansion^92^, which is the most common form of genetic ALS and FTLD^93^. These findings point to a key role of lysosomal stability and function in governing pathological TDP-43 aggregation. In view of these important links between lysosomal (dys)function and neurodegeneration, and to understand if the TMEM106B-mediated increase in seeding potency mimics lysosomal disruption, we used an independent approach to injure the lysosomes and assessed the effects on seeding outcomes. To that end, we used the lysosomotropic agent LLME, which accumulates specifically in lysosomes, where it turns into membrane-disruptive oligomers, causing lysosome permeabilization^70^. Combining LLME with concentrated recombinant LCD fibrils resulted in a marked increase (∼50-70 times more) of the seeding potency in human neurons. In fact, we observed that in the absence of LLME, the majority of internalized LCD fibrils remained confined to the lysosomal compartment, where they were neutralized. Conversely, FTLD-TDP-A seeds showed 20 to 100 times higher seeding efficiency compared to LCD fibrils even in the absence of LLME-induced lysosomal injury. Notably, this seeding activity could be ranked according to their TMEM106B fibril content. This strong correlation between TMEM106B fibril load and seeding outcomes suggests that TMEM106B fibrils may compromise lysosomal integrity, facilitating TDP-43 fibril escape and enhancing seeding. It is conceivable that TMEM106B fibrils within patient extracts seed fibrilization of endogenous TMEM106B within lysosomes, thereby driving lysosomal rupture and potentiating TDP-43 aggregation and loss of function.

Recently, the effect of compartmentalization and lysosomal escape on the seeding activity of internalized tau fibrils has been described^94, 95^. In line with our data on TDP-43, cells with pre-existing lysosome instability were seeded by tau fibrils at a higher rate than the general population^95^, suggesting that lysosomal escape might be a common mechanism enhancing propagation of intracellular aggregates in multiple neurodegenerative diseases.

While seeded human neurons developed both intranuclear and cytoplasmic TDP-43 pathology, LLME treatment preferentially amplified the cytoplasmic aggregates. This observation suggests that these two pathological phenotypes, which are well-documented in ALS/FTLD patient tissues^1, 2, 72–74^, may arise through distinct mechanistic pathways. Given that growth-arrested SH-SY5Y cells did not develop intranuclear TDP-43 inclusions in either the current study or our previous report^44^, and because seeding occurred during neuronal differentiation, we cannot rule out a potential contribution of cell division to intranuclear aggregate formation. Cell division might reduce the nuclear protection naturally exerted by the nuclear membrane, thus favoring the penetration of exogenous aggregates. However, the description of intranuclear TDP-43 inclusions in non-dividing neurons of postmortem human tissue^1, 2, 72–74^ indicates that such aggregates can arise independently of cell division.

FTLD-TDP-A patient extracts induced markedly stronger neoaggregation, loss of function and toxicity than either FTLD-TDP-C extracts or recombinant LCD fibrils, in line with earlier studies^5, 18–20^. This divergence indicates that factors beyond TDP-43 fibrils shape seeding and downstream toxicity and our data identify TMEM106B fibrils as key contributors potentiating the effects of FTLD-TDP-A seeds. Additional, yet unidentified components and/or structural fibril features may play important roles. ANXA11, which forms heterotypic fibrils together with TDP-43 specifically in FTLD-TDP-C, is an excellent candidate that may influence seeding outcomes in this subtype. Our analysis showed that while ANXA11 fibrils are specific to FTLD-TDP-C seeds, as previously reported^18, 28, 29^ their levels do not influence seeding. In fact, FTLD-TDP-C seeding effects were overall weak and did not reach statistical significance, albeit we still detected some neoaggregates and occasional cells with TDP-43 loss of function. However, the accompanying mScarlet intensity, reflecting nuclear TDP-43 loss, was modest and endogenous TDP-43 recruitment was comparatively weak in both neuron-like cells and neurons. Future work will investigate whether the levels of ANXA11 in the recipient cells might influence seeding outcomes of FTLD-TDP-C fibrils.

In conclusion, our study provides quantitative evidence for distinct molecular mechanisms underlying FTLD subtype pathology through patient-derived seeding models. We identified pathological TDP-43 and TMEM106B fibrils as key factors driving seeding capacity and subsequent loss of function in neuron-like cells and human neurons and revealed time-dependent changes in TDP-43 interactome that distinguish FTLD subtypes. Importantly, we demonstrate that lysosomal rupture strikingly potentiates the induction of TDP-43 aggregation and loss of function in the presence of pathological seeds in human neurons. Our findings underscore a central role for lysosomal dysfunction in the development of TDP-43 pathology and establish the most robust model to date for seeded aggregation and loss of function. We anticipate that this model will be instrumental for identifying modifiers and therapeutic targets for TDP-43 proteinopathies.

## Acknowledgments

We are grateful to all the patients and their families for donating tissues for scientific research. This work would not have been possible without their generosity and foresightedness. We gratefully acknowledge the Netherlands Brain Bank, Netherlands Institute for Neuroscience, Amsterdam for postmortem brain tissues. This study was supported by the Swiss National Science Foundation (Project Grant 320030-231921) and by the Association for Frontotemporal Dementia (FTD Biomarkers Initiative Grant) to M.P., a Live Like Lou postdoctoral fellowship to C.S., a Byrne Family and Judith & Pape Adams Fellowship and a Gilda Slifka Neuroscience Transformative award to N.R., an ALS Scholar in Therapeutics award from The Healey Center at Mass General and ALS Finding a Cure to S.M.L, the Chan Zuckerberg Initiative to C.L-T and M.W., and the NIH/NINDS (RM1NS133601 to M.W. and C.L-T). The authors would like to acknowledge the Functional Genomics Center Zurich (FGCZ) for the training and technical support in all the proteomics experiments; the cytometry facility and the center for microscopy and image analysis (ZMB) at University of Zurich for the technical help with FACS experiments, microscopy and image data handling, respectively. We thank all the members of the Polymenidou lab for the critical feedback and discussions. All schematics included in figures have been generated using BioRender.

## Author contributions

W.Z., CS and M.P. conceptualized the study, designed the experiments and wrote the manuscript. W.Z. performed and analyzed the patient sample characterization, seeding, quantifications and proximity proteomics experiments in SH-SY5Y cells. CS performed and analyzed the seeding, FACS and imaging experiments in *NGN2* neurons and devised the lysosomal rupture paradigm. B.G. generated the TDP-REG *NGN2* iPSC line and devised the protocol for neuron dissociation for FACS analyses. B.G. and C.S. developed together the seeding conditions with *NGN2* neurons and performed the seeding experiments. M.H. helped in performing all seeding experiments with neurons, the respective FACS analyses and in figure building and revision. N.L. helped in the FACS analyses of seeded neurons, in image acquisition and performed all image-based manual quantifications. N.L. also helped in figure building and revision. F.L. helped in sample preparation for interaction proteomics experiments and in figure building and revision. F.L. also performed part of the image-based quantifications on seeded neurons. M. Peter helped in processing patient postmortem brain samples. C. L-T., M.W. and N.R. supervised the generation of the TDP-REG *NGN2* iPSC line. S.M.L. generated the iPSC lines expressing the NGN2 constructs used in this work. B.R. provided training and feedback in performing LC/MS analysis. S.J. produced and characterized the TDP-43 LCD recombinant fibrils. R.M. produced and characterized the phosphorylated recombinant TDP-43 used for loading control on western blots. E.D.C. and A.A. provided the tau and α-synuclein fibrils. O.W. and P.F. invented and characterized the TDP-REG system. L.P. suggested the initial TMEM106B analyses on patient samples and provided the fibrillar core-specific antibody. M.P. directed the entire study. All authors read and edited the manuscript.

## Material and Methods

### Postmortem brain tissue

Patient samples were kindly provided by the Netherlands Brain Bank (NBB), Netherlands Institute for Neuroscience, Amsterdam (**Table S3**).

### SarkoSpin preparation of patient material

Homogenization of patient brain tissue was performed as previously described^18, 63^. After homogenization of patient brain tissue, 150 µL of brain homogenate (contains approximately 30 mg of frontal cortex per aliquot) were prepared in high salt (HS) buffer 10 mM Tris-HCI (Sigma-Aldrich, T3253-1KG), pH 7.4, 150 mM NaCl (Sigma-Aldrich, 71380), 0.5 mM EDTA (Amresco, 0105-1KG), 1 mM DTT (Sigma-Aldrich, R0861), cOmplete mini EDTA-free protease inhibitors (Roche, 11836170001), PhosphoSTOP phosphatase inhibitors (Roche, 4906837001), and stored at -80°C. Subsequently, 50 µL of HS buffer with benzonase mix (Merck Millipore, 70664) and 14 mM MgCl_2_ (Sigma-Aldrich, M8266-100G) was added to each aliquot to digest nucleic acids for 5 min at RT. Following this, 200 µL of 2X HS buffer containing 4% w/v N-lauroyl-sarcosine (Sigma-Aldrich, L5125-100G) was added, and samples were incubated on a thermoblock for 45 min at 38°C with 600 rpm shaking to solubilize non-aggregated proteins. After the solubilization, 400 µL of ice-cold HS buffer with 40% sucrose (Sigma-Aldrich, 84100) was added and the samples were centrifuged at 21’200 x g for 30 min at RT to pellet the sarkosyl-insoluble protein fraction. The supernatant was removed, and pellets were washed twice with 200 µL of 1X PBS to remove as much of remaining lipids and sarkosyl as possible. Pellets for immediate use were resuspended in 100 µL of PBS by sonication (QSonica Q500 sonicator, amplitude = 60%, pulse = 2 s on/2 s off, total 3 min), or snap frozen on dry ice and stored at -20°C prior to use.

### Fluorescent labeling of patient-derived material

For one SarkoSpin (SKS) pellet derived from 30 mg of brain tissue to 20 parts of soluble protein, 1 part of 0.2 M sodium bicarbonate (Sigma-Aldrich, S6014) solution adjusted to pH 9.0 with 2 M sodium hydroxide (Sigma-Aldrich, S5881), was added to obtain a pH of 8.3. 5 nmol of Atto-488 NHS-ester (ATTO-TEC, AD 488-31) resuspended in DMSO (Sigma-Aldrich, D2650) was added to the solution and incubated overnight at 4°C. The labeled SKS pellets were collected by centrifugation at 21’200 x g at RT for 20 min and washed with PBS twice by gentle vortexing and centrifugation at 21’200 x g at RT. Finally, the SKS pellets were resuspended in PBS and sonicated with the program described above prior to use for seeding experiments.

### Production of recombinant TDP-43 LCD, α-synuclein and tau K18 fibrils

After purification of the recombinant TDP-43 low complexity domain fragment, unlabeled and Atto-488 labeled fibrils were generated as previously described^44^. The Alexafluor-488 α-synuclein and tau K18 fibrils were generated and labeled as previously described in Scialò et al.,^44^ (α-synuclein and TDP-43 LCD fibrils), and De Cecco et al.,^69^ (tau K18 fibrils).

### SDS-PAGE and western blotting

For chemiluminescent western blot of BioID2-TDP-43WT, BioID2-TDP-43mNLS and BioID2-GFP cell lysates, SH-SY5Y cells were plated in 6-well plates at a density of 0.3 x 10^6^ cells per well. When cells reached 70% confluency, usually the day after plating, they were growth arrested with biotin-depleted medium containing BioLock (IBA Lifesciences, 2-0205-050) at 2 µl/ml for 48 hours. The medium was then replaced with fresh medium supplemented with biotin (Sigma-Aldrich, B4501) to a final concentration of 50 μM for 24 hours. Post incubation with biotin, cells were washed twice with PBS and lysed in RIPA buffer (ThermoFisher, 89900) supplemented with 1mM DTT (ThermoFisher, R0861), 1.5 mM MgCl_2_ (Thermo Fisher, AM9530G) and 1 tablet of cOmplete mini EDTA-free protease inhibitors (Roche, 11836170001) per 10 ml of RIPA buffer. Per well of a 6-well plate, 0.5 μl of benzonase mix (Merck Millipore, 70664) was added and the lysate was incubated for 5 minutes. Protein amount in the samples was measured using a BCA assay, according to the manufacturer’s instructions (ThermoFisher, 23225). For chemiluminescent western blot of patient samples, 10% of the sarkosyl-insoluble protein was loaded per lane, which approximately corresponds to 17 μg of protein.

All reagents used are from ThermoFisher unless otherwise specified. Protein samples were mixed with 1X LDS loading buffer and 1X Bolt sample reducing agent (Life Technologies, B0009), denatured at 70 °C for 10 min, and loaded onto NuPAGE 4-12% BisTris Plus Gels. Proteins were transferred from the gels onto nitrocellulose membranes using iBlot 2 Transfer NC Stacks with iBlot 2 Dry Blotting System (protocol: 7 min at 20 mV). Membranes were blocked with EveryBlot blocking buffer (Bio-Rad, 12010020) for 20 minutes and incubated with primary antibodies (Anti-TDP-43 3H8, Novus Biologials, NBP1-92695, 1:1000; TDP-43 N-term and C-term cocktail, Proteintech, 10782-2 and 12892-1, 1:2500; Anti-HA, Cell Signaling, 3824S, 1:1000; TDP-43 pS409/410, CosmoBio, TIP-PTD-M01, 1:2500; gift antibody against TMEM106B fibrillar core provided by Petrucelli lab, 1:2000) in EveryBlot blocking buffer overnight at 4 °C. After this, membranes were washed three times with PBST, followed by incubation with HRP-conjugated secondary antibodies (HRP Conjugated AffiniPure Goat anti-mouse IgG, Jackson ImmunoResearch, 115-035-146, 1:5000; HRP Conjugated AffiniPure Goat anti-rabbit IgG, Jackson ImmunoResearch, 115-035-144, 1:10000; Streptavidin-HRP, Abcam, ab7403, 1: 40000) in EveryBlot blocking buffer for 3 h at room temperature. After three washes in PBST, immunoreactivity was visualized by chemiluminescence using SuperSignal West Pico or Femto Chemiluminescent Substrate (ThermoFisher, 34577, 34096) on a Fusion FX6 EDGE imager (Vilber).

### Human neuroblastoma SH-SY5Y cell culture

Human neuroblastoma SH-SY5Y cells (ECACC, 94030304) were cultured for up to 20 passages in Dulbecco’s Modified Eagle Medium/Nutrient Mixture F-12 (DMEM/F-12) (Gibco, 21331046) supplemented with 10% fetal bovine serum (FBS) (Gibco, A5256701) and 1% Penicillin-Streptomycin (Gibco, 15140122) at 37°C in a humidified atmosphere with 5% CO_2_. Cells were regularly tested for mycoplasma contamination.

### Generation of the BioID cell lines

The plasmid for mammalian expression of BioID2 was purchased from Addgene (74224) and modified using the NEBuilder HiFi DNA Assembly Cloning Kit (NEB, E5520) to insert the sequences of TDP-43 or GFP. The Q5 Site-Directed Mutagenesis Kit (NEB, E0554S) was used to mutagenize the nuclear localization signal sequence of TDP-43 as previously described^76^. BioID2-TDP-43WT/mNLS or BioID2-GFP constructs were packaged into lentivirus as previously described^17^. In brief, HEK293T cells (ATCC, CRL-3216; gift from the Greber laboratory) were adapted to serum-free OHN medium and co-transfected with Lipofectamine 2000 (Invitrogen, 11668019; 8.3 µl/ml final volume) and the CMV-Gag-Pol (Harvard, dR8.91) and pVSV-G (Clontech, part of 631530) plasmids. OHN medium was based on Opti-MEM (ThermoFisher, 11058-021) supplemented with 0.5% B27 (ThermoFisher. 12587-010), 0.5% N2 (ThermoFisher, 175020-01), 1% GlutaMAX (ThermoFisher, 35050038), and 25 ng/ml bFGF (ThermoFisher, PHG0261). Serum-free conditions were adopted to reduce basal expression from the transfer vector (that is, it eliminates traces of tetracyclines in the FBS) and prevent serum carryover into the viral supernatant. Medium was changed the following morning, and viral supernatants were collected 48 h post-transfection (36 h after medium change), centrifuged (500g, 10 min, 4 °C), filtered through a Whatman 0.45-µm CA filter (GE, 10462100), and concentrated overnight using Lenti-X Concentrator (Takara, 631232) according to the manufacturer’s instructions. The resulting lentiviral pellets were resuspended in culture medium to achieve 10X concentrated preparations, which were titrated using Lenti-X GoStix Plus (Takara, 631280). All lentiviruses were used at 500 ng (GV)/ml of medium in a total volume of 1 ml in sub-cultured 6-well plates. Medium was completely exchanged the following day to medium containing 1200 µg/ml Geneticin (Invivogen, ant-gn-5), and cells were maintained as stable lines thereafter.

### Seeding on SH-SY5Y cells

After coating with poly-D-lysine (Sigma-Aldrich, P6407) for 1 h and washing three times with sterile ddH_2_O, SH-SY5Y cells were plated at 0.1 x 10^5^ cells per well in a µ-Plate 96 Well Square (Ibidi, 89627), 0.5 x 10^5^ cells per well in a 24-well plate, or 0.3 x 10^6^ cells per well in a 6-well plate. When cells reached 70% confluency, usually the day after, they were growth arrested in biotin-depleted medium containing BioLock (IBA Lifesciences, 2-0205-050) at 2 µl/ml and maintained arrested throughout the experiment. One day later, cells were inoculated with freshly sonicated patient-derived SarkoSpin extracts at a dosage of 2.5% of the total resuspended and sonicated pellet/cm^2^ surface area of the treated well. Medium was changed every 3 days until the end of the experiment.

### FACS-based quantification of internalized labeled seeds

BioID2-TDP-43WT or mNLS stable cells were plated in 24-well plates at a density of 0.5 x 10^6^ cells per well and seeded as described above. Before flow cytometry analysis, cells were trypsinized and resuspended in FACS buffer (1% FBS (Gibco, A5256701), 25 mM HEPES (Sigma-Aldrich, H3537), and 5 mM EDTA (Sigma-Aldrich, E4884) in 1X PBS (Gibco, 10010023)). The cell suspension was filtered through a 35 µM cell-strainer-coupled Falcon tube (Corning, 352235) to obtain a single-cell suspension before loading onto a BD FACSCanto II flow cytometer (BD Biosciences). After gating for singlet cells, the Atto488+ gate was defined as the top 99.99% quantile of fluorescence intensity in the no treatment (mock) condition x 1.05.

### Generation of iPSC-derived neuronal culture carrying TDP-REG reporter

The parental KOLF2.1J iPSC line (iNDi initiative)^96^ had been previously engineered to express, under a doxycycline-inducible promoter, the human Neurogenin-2 (*NGN2*) delivered via Piggybac plasmid^68^. The plasmids for the expression of the TDP-REG system and for the hyperactive piggyBac transposase as well as their usage for generation of a polyclonal SH-SY5Y stably expressing TDP-REG were previously characterized^44^. In order to introduce the TDP-REG reporter for the generation of a polyclonal iPSC line, cells were plated onto T75 flasks (Thermo Scientific, 156800) coated with 5ug/mL Vitronectin (Stem Cell Technologies, 100-0763) in StemFlex Medium (Gibco Life Technologies, A3349401) and 1X RevitaCell (Gibco Life Technologies, A26445-01). One day after thawing, complete media change without RevitaCell was performed and iPSCs were maintained at 37 °C with 5% CO_2_ with daily media change until passaging. Upon reaching 80% confluence, cells were lifted using Accutase (Stemcell Technologies, 07920) and 1 x 10^6^ cells per condition were subjected to nucleofection (Human Stem Cell Nucleofector Kit 2, Lonza, VPH-5022; Nucleofector 2b Device, Lonza). Nucleofection was performed using the piggybac TDP-REG vector together with a vector expressing the hyperactive piggyBac transposase at a 1:1.25 ratio of transposase vector to TDP-REG vector. Upon nucleofection, cells were re-plated in StemFlex medium with 1X Revitacell in one well of a 12-well plate (Thermo Scientific, 150200). Selection was performed for at least 15 days in StemFlex medium containing 0.2 mg/mL blasticidin (Invivogen, ant-bl-05). To verify the efficiency of selection, a control transfection without the transposase expression vector was carried out in parallel.

### Cloning and Lentiviral (LV) production

The lentiviral transfect vector for the knockdown of TARDBP was generated as previously described^44^. Briefly, the pSHE LV vector was modified using the Q5 Site-Directed Mutagenesis Kit (NEB, E0554S) to insert either *TARDBP* or control shRNA under a hU6-driven promoter, and the NEBuilder HiFi DNA Assembly Kit (NEB, E5520S) to replace EGFP with HaloTag downstream of an inducible mTRE promoter. The lentiviral transfect vector was generated, harvested and concentrated as described previously^17^. Lentiviral pellet was resuspended in N Dulbecco’s Modified Eagle Medium:Nutrient Mixture F-12 (DMEM/F-12, Gibco, 31330038) to obtain 10X concentrated preparations. To determine the optimal viral dose, a viral titration was performed in SH-SY5Y cells (ECACC, 94030304), where cells were transduced with increasing volumes of concentrated viral supernatant. The minimal volume achieving >80% transduction efficiency without cytotoxicity was used for subsequent experiment.

### Neuronal differentiation and lentiviral transduction

Neuronal differentiation was started by plating iPSCs onto Matrigel-coated (Corning, 354230) 6-well plates at a density of 0.5x10^6^ cells per well in induction medium supplemented with 2 µg/mL doxycycline (Sigma, D9891-1g) and 1X RevitaCell (Gibco, #A2644501). Induction medium consisted of Dulbecco’s Modified Eagle Medium:Nutrient Mixture F-12 (DMEM/F-12, Gibco, 31330038) complemented with 1X Glutamax (Gibco, 35050061), 1X Non-essential amino acids (Corning, 25-025-Cl) and 1X N-2 Supplement (Gibco, 17502048). For the expression of the psHalo vector carrying TARDBP shRNA, cells were transduced immediately after plating with the optimized viral dose to reach >80% of transduction efficiency (equivalent to 30 µL viral stock mixed in 1.5 mL medium for 0.5 x 10^6^ cells plated in a well of the 6-well plate). After 24 hours, a complete medium change without RevitaCell was performed to remove residual viral particles, while on day 2 only a half-medium change was performed. At day 3 of differentiation, cells were replated onto 24-well plates at a density of 0.5 x 10^6^ cells per well and the medium was switched to *NGN2* differentiation medium consisting of Neurobasal medium (Gibco, 21103049) complemented with 1X Glutamax (Gibco, 35050061), 1X B-27 Supplement (Gibco, 17504044) and 1% of a 5M NaCl solution prepared by dissolving NaCl (Sigma Aldrich, 17380) in Ultra-Pure Distilled Water (Invitrogen 1097749). Prior to usage *NGN2* differentiation medium was freshly supplemented with 2 µg/mL doxycycline, 5 µM aphidicolin (Cell Signaling Technology, 32774), 10 ng/mL BDNF (Lifetech, PHC7074) and 10 ng/mL NT3 (Peprotech, 450-03). Half medium changes were performed every other day until day 7 of differentiation, when cells were dissociated and analyzed by Flow cytometry (FACS).

### Seeding of TDP-REG *NGN2* neurons

TDP-REG iPSCs were directly plated on day 0 of differentiation on Dendritic Polyglycerol Amine (dPGA) (Dendrotek Biosciences Inc-PO) and Matrigel-coated (Corning, 354230) 96-well (IBIDI) (at a density of 0.15x10^5^) or 24-well plates (TPP) (at a density of 0.2 x 10^6^) in induction medium supplemented with 2 µg/mL doxycycline (Sigma, D9891-1g) and 1X RevitaCell (Gibco, #A2644501). After 24 h, fresh induction medium supplemented with doxycycline was mixed with sonicated in vitro generated fibrils (at a final concentration of 200 nM) or patient-derived SarkoSpin extracts (dosage of 5.3% of the total resuspended and sonicated pellet/cm^2^ surface area of the treated well) and added to differentiating neuronal progenitor cells (NPC) for 2 days to allow uptake. On day 3 of differentiation, induction medium was replaced with *NGN2* differentiation medium supplemented with 2 µg/mL doxycycline, 5 µM aphidicolin (Cell Signaling Technology, 32774), 10 ng/mL BDNF (Lifetech, PHC7074) and 10 ng/mL NT3 (Peprotech, 450-03). Half medium changes were performed every other day until day 7 of differentiation, when cells were dissociated and analyzed by Flow cytometry (FACS) or fixed for IF. When performed, incubation for 30 minutes with 1 mM L-leucyl-L-leucine, methyl ester LLME (Cayman Chemical) was performed on day 3 of differentiation after the 2 days of incubation with fibrils or SarkoSpin extracts and before medium change with supplementedifferentiation medium.

### Cell dissociation and FACS-based quantification of TDP-REG reporter activation in *NGN2* neuronal culture

For all FACS-based experiments in *NGN2* neurons the same procedure was applied. *NGN2* neurons were washed once with PBS (Gibco, 70011044) and dissociated with a Papain dissociation kit (Worthington Biochemical, LK003150) prepared according to manufacturer’s instructions. For each well of the 24-well plate, 300 µL of papain solution and DNase were added and cells were incubated at 37 °C with 5% CO_2_ for 5 minutes. Following addition of 300 µL of *NGN2* medium supplemented with all compounds and 1X RevitaCell (Gibco, A2644501) for each well, the cell monolayer was gently dislodged and mixed to obtain a single cell suspension. Cells were centrifuged at 200 x g for 5 minutes and resuspended in 200 µL FACS Buffer (5 mM EDTA (Sigma-Aldrich) in PBS 1X (Gibco). Cells were filtered through a 35 µm strainer-coupled Falcon (Corning, 352235) to obtain a single-cell suspension and samples were analyzed using a BD FACSymphony flow cytometry instrument (BD Biosciences).

### Immunofluorescence

SH-SY5Y cells were fixed with formaldehyde (Pierce, 28908) added directly to cell culture medium at a final concentration of 4% for 20 minutes at room temperature. After three washes with PBS (Gibco, 10010023), cells were permeabilized in a permeabilization buffer composed of 50 mM NH_4_Cl (Sigma-Aldrich, 254134-5G) with 0.25% Triton (Sigma-Aldrich, X100-1L) in PBS for 1 hour at RT. After 2 washes with PBS, cells were then incubated with the primary antibodies (Anti-TDP-43 Clone, 671834 aa 1-103, Novus Biologicals, MAB7778, 1:500; Anti-HA, Cell Signaling 3824S, 1:1000) in a blocking solution composed of 10% donkey serum (Sigma-Aldrich, S30-M), 3% BSA (Sigma-Aldrich, A9418-50G), 0.25% Triton in PBS overnight at 4°C. After three washes with PBS, cells were subsequently incubated with fluorescently labelled secondary antibody (Alexa Fluor 647 Donkey anti-Rabbit IgG, ThermoFisher, A-31573; Alexa Fluor 647 Donkey anti-Mouse IgG, ThermoFisher, A-31571) diluted at 1:1000 in blocking solution for 3 hours at RT, followed by three washes with PBS. After removing wash buffer from the cells, 3 drops of mounting medium with DAPI (ibidi, 50011) were added to the well and the cells were incubated for 4 hours prior to imaging or storage at 4°C. TDP-REG *NGN2* neurons were fixed for 20 minutes at RT by direct addition of 4% formaldehyde (Pierce, 28908) and 0.05% glutaraldehyde solution (3802-75mL Sigma Aldrich) to culture medium. After three washes with PBS, cells were permeabilized for 5 minutes at room temperature in permeabilization buffer consisting of 0.1% Triton (Sigma-Aldrich, X100-1L) in PBS. Neurons were then incubated 30 minutes at room temperature in blocking buffer composed of 10% donkey serum (Sigma-Aldrich, S30-M) and 0.1% Triton in PBS followed by incubation with primary antibodies (Anti-TDP-43 Clone 671834 aa 1-103, Novus Biologicals, MAB7778, 1:500; Anti-MAP2, Abcam, ab5392, 1:1000; Anti-FUS, Thermo Scientific, A300-293A, 1:000; Anti-p62, Proteintech, 18420-1-AP, 1:500; Anti-LIMP2, Novus biologicals, NB400-129, 1:200) diluted in blocking buffer for 2 hours at room temperature. After 3 washes with PBS, cells were incubated with secondary antibodies (Goat anti-Chicken IgY (H+L) Alexa Fluor Plus 405, Thermo Fisher, A48260, 1:1000; Donkey anti-Rabbit IgG (H+L) Alexa Fluor 488, Thermo Fisher, A-21206; Donkey anti-Mouse IgG (H+L) Alexa Fluor 647, Thermo Fisher, A-31571; Donkey anti-Rabbit IgG (H+L) Alexa Fluor 647, Thermo Scientific, A-31573) diluted 1:1000 in blocking buffer for 1 hour at room temperature. DAPI (ibidi, 50011) was diluted 1:1000 in permeabilization buffer and incubated for 5 minutes at room temperature.

### Quantification of colocalizing TDP-43 aggregates

This paragraph refers to quantifications shown in Figures **2h-i** and **S4b, c.** For the quantification of neo-aggregate volume, defined as colocalized endogenous TDP-43-HA signal with exogenous fluorescently labeled SKS aggregates, images were taken with an ImageXpress Confocal HT.ai High-Content imaging system (Molecular Devices) maintained by the Center for Microscopy and Image Analysis (ZMB) at the University of Zurich. 16-bit images were acquired using a Nikon LWD Lambda S 40XC water-immersion objective with a 2048 x 2048 image size, corresponding to 0.168 µm per pixel in xy. Z-stacks were collected over a 5.8 µm depth with a z-step size of 0.2 µm. 6 fields of view (FOV), containing an average of 457 cells, were imaged per condition with the microscopic and image processing parameters kept constant between repetitions. All images were deconvolved with SVI Huygens Professional version 24.04, with the signal-to-noise ratio set to 42.5 for DAPI, 44.9 for Atto-488 and 37.4 for HA. Subsequently the images were processed with Imaris Version 10.2.0 by reconstructing the image using the ‘‘Surfaces’’ function. Nuclear surfaces were created from the DAPI channel, with a surface detail of 0.3 µm from pixels with an automatic intensity threshold. Objects with a volume of less than 50 µm^3^ were excluded. To separate individual nuclei, touching objects were split under intensity-based mode with a seeding diameter of 9 μm. Surfaces for exogenous aggregates were created from the Atto-488 channel with the surface detail set to 0.01 μm from pixels with an intensity over 400. Surfaces for TDP-43-HA were created from the HA channel, after applying a 3x3x3 pixel median filter to remove salt-and-pepper background noise, with the surface detail set to 0.1 μm from pixels with an intensity threshold set over 100. To separate punctate TDP-43-HA signal, touching objects were split under intensity-based mode with a seeding diameter of 1 μm. HA-surfaces that had an overlapping volume of more than 30% with an exogenous aggregate object were counted as neo-aggregates. Objects with a volume of less than 0.003 μm^3^ were excluded. Statistical data on the volume of each surface object were exported to calculate the total volume of neo-aggregates, normalized to the count of DAPI surfaces in an FOV, for comparing the differences between the conditions.

### High-content imaging and quantification on TDP-REG cells

This paragraph refers to quantifications shown in Figures **4b-e**. Image acquisition was performed on an ImageXpress Confocal HT.ai High-Content imaging system (Molecular Devices) maintained by the Center for Microscopy and Image Analysis (ZMB) at the University of Zurich. 16-bit images were acquired using a Nikon CFI Plan Apo Lambda 10X/NA 0.45 objective with a 2048 x 2048 image size, corresponding to 0.678 mm per pixel in xy. Z-stacks were collected over a 33.9 mm depth with a z-step size of 3.36 mm. 25 fields of view (FOV), covering 90% of the bottom area of the treated well, were taken per condition. All images were stitched into stacks using the MD2Hyperstack script developed by the ZMB. The stitched images were converted to 2D by summing the intensity over the slices, and the nuclear segmentation was performed by implementing *StarDist* on the DAPI channel^97^. After subtracting the background by division from the autofluorescent channel for the mScarlet channel, the processed mScarlet intensities and TDP-43 intensities within each segmented cell nucleus were exported. Objects with TDP-43 intensities higher than 3 standard deviations were excluded, since such high intensity usually indicates artefacts instead of real cells. TDP-REG positive cells were selected by filtering the list of all objects using a thereshold defined as the top 99.99% quantile intensity in the no treatment condition x 1.05. Our filtering and plotting utilities are open-sourced together with the data we generated (code available on: https://github.com/Chocobosaurus/TDPLoF).

### Quantification of the localization of p62/TDP-43-positive aggregates in TDP-REG *NGN2* human neurons

This paragraph refers to image quantifications shown in **Fig. 6m**, **Fig. 7e**, **Fig. S10f and Fig. S11d**. Image acquisition was performed on a Leica SP8 inverse Falcon confocal microscope equipped with an HC PL APO corr. CS2 63 x/Nam1.4 oil-immersion objective. Images were collected at 512 x 512 resolution in 8-bit depth, and all microscopic and image processing parameters were kept constant across repetitions. For each condition, 10-12 fields of view (FOVs) were acquired, each containing at least one mScarlet-positive neuron. Z-stacks were collected for each FOV using 0.5µm optical step size (19 slices). All images were deconvolved with SVI Huygens Professional version 24.04 with the signal to noise ratio set as 42.5 for channel DAPI, 44.9 for Atto488 and 37.4 for HA, respectively. Subsequently the images were processed with Imaris Version 10.2.0 by reconstructing the image using the ‘‘Surfaces’’ function. Nuclear surfaces were created from the DAPI channel, with a surface detail of 0.3 µm from pixels with an automatic intensity threshold. Objects with a volume of less than 250 µm^3^ were excluded. To separate individual nuclei, touching objects were split under intensity-based mode with a seeding diameter of 9 μm. Surfaces were created from the p62 channel after applying a 3x3x3 pixel size median filter for removing salt and pepper noise in the background, as well as a step of baseline subtraction with a parameter set at 0.5. The surface detail set to 0.1 μm from pixels with an intensity over 3.5. Objects with a volume of less than 0.03µm^3^ were excluded. Surfaces for TDP-43 were created after applying a 3x3x3 pixel size median filter for removing salt and pepper noise in the background, with the surface detail set to 0.1 μm from pixels with an intensity threshold set over 6. To separate punctate TDP-43 signal, touching objects were split under intensity-based mode with a seeding diameter of 2 μm. Such objects that have an overlapping volume of over 30% with an p62 object are counted as aggregates. Objects with a volume of less than 0.003 μm3 were excluded. Statistic data on the volume of each surface object was exported for calculating the overlap ratio with DAPI surfaces. Objects with a value > 50% were counted as localized in nuclear regions.

### Manual quantification of LIMP2 retained LCD-488 and patient-derived SKS aggregates in TDP-REG *NGN2* human neurons

This paragraph refers to image quantifications shown in **Fig. 6g-h** and **Fig. 7g**. Image acquisition was performed on a Leica SP8 inverse Falcon confocal microscope equipped with an HC PL APO corr. CS2 63 x/Nam1.4 oil-immersion objective. Images were collected at 512 x 512 resolution in 8-bit depth, and all microscopic and image processing parameters were kept constant across repetitions. For each condition, 10-12 fields of view (FOVs) were acquired, each containing at least one mScarlet-positive neuron. Z-stacks were collected for each FOV using 0.5µm optical step size (19 slices), and full stacks were examined during all manual quantifications. The quantification of the retained and escaped exogenous TDP-43 LCD-488 aggregates shown in **Fig. 6g-h** was performed manually in Fiji. For each FOV, one mScarlet-positive and one negative neuron were analyzed, and all LCD-positive inclusions were counted along with the subset colocalizing with LIMP2, indicating lysosomal sequestration. Colocalization was assessed by manual slice-by-slice inspection of the z-stacks. The quantification of autofluorescent SKS aggregates shown in **Fig. 7g** was performed manually in Fiji. All mScarlet-positive neurons per FOV were examined for the presence of SKS material and scored for colocalization with endogenous TDP-43 (indicating a recruiter) and with LIMP2 (indicating lysosomal sequestration) by manual slice-by-slice inspection of the z-stacks. In the CTRL patient sample, only negative neurons were analyzed, and only non-recruiting SKS aggregates were present and quantified for their colocalization with LIMP2. An example illustrating the criteria used for these manual assessments is shown in **Fig. S11e**.

### Image-based quantification of TDP-43 and FUS nuclear intensity in TDP-REG *NGN2* human neurons

This paragraph refers to image quantifications shown in **Fig. S10 c and d**. Image acquisition was performed on a Leica SP8 inverse Falcon confocal microscope equipped with an HC PL APO corr. CS2 63 x/Nam1.4 oil-immersion objective. Images were collected at 512x512 resolution with 8-bit depth, and all imaging and image processing settings were kept constant across repetitions. For each condition, 10-12 fields of view (FOVs) were acquired, each containing at least one mScarlet-positive neuron. Z-stacks were collected for each FOV using 0.5 µm optical step size (19 slices), and full stacks were examined during all quantifications. Nuclear intensities have been calculated using a dedicated Macro in Fiji. Briefly, the macro generates two ROIs, one from the mask for DAPI signal within mScarlet neurons (nuclei of mScarlet neurons) and another for DAPI outside mScarlet neurons (all other DAPI-positive nuclei), Finally, the macro measures the mean intensity of TDP-43 and FUS within these two ROIs.

### Proteomics analysis Sample preparation

SH-SY5Y cells were plated on 6-well plates (TPP) at a density of 3 x 10^5^ cells per well. Upon reaching 70% confluency, typically the day after, cells were growth arrested by biotin-depletion using biotin blocking solution (Biolock, iba lifescience, 2-0205-050) added to the cell culture medium at 2 µl/ml. Cells were maintained in growth arrested conditions for 48 hours for the identification of nuclear and cytoplasmic TDP-43 interactors using BioID2-TDP-43WT, BioID2-TDP-43mNLS and BioID2-GFP cell lines, respectively, or until one day before the endpoint of the experiment for the identification of the TDP-43 interactome with patient material seeding. Subsequently, the medium was replaced and supplemented with biotin (Sigma-Aldrich, B4501) to the final concentration of 50 μM for 24 hours. Post incubation with biotin, cells were washed twice with PBS and directly lysed in a lysis buffer, which was composed of 8M Urea, 1mM DTT (Thermo Fisher, R0861), 1.5mM MgCl2 (Thermo Fisher, AM9530G) with 1% Triton (Sigma-Aldrich, X100-1L) in 50mM Tris buffer, pH 7.4. This harsh lysis step was performed to disrupt all biological assemblies and release the biotinylated proteins. For each well of a 6-well plate, 150 μl of lysis buffer supplemented with 0.5 μl of benzonase were added and the lysate was incubated for 5 minutes. Afterwards, the lysate was sonicated with a QSonica Q500 sonicator at amplitude = 60%, pulse = 2 sec. on/2 sec. off for a total time of 3 minutes, followed by a centrifugation step at 16500 x g for 10 minutes to remove debris and remaining DNA in the sample. The protein amount in the supernatant was measured using a BCA assay for normalizing the input for affinity enrichment of the biotinylated proteins. Per sample, lysates form four wells of a 6-well plate were pooled and incubated with 4 μl of Streptavidin Sepharose beads (Cytiva, 17511301). Streptavidin Sepharose beads were precleared twice in 50mM Tris buffer and once in lysis buffer by shaking on a thermomixer (Eppendorf) at 800 rpm at 27°C for 7 minutes followed by centrifugation at 4000 x g for 2 minutes. The input was then added to the beads, followed by incubation of 6.5 hours at room temperature with rotation. After the incubation, the supernatant was removed and the beads were resuspended in 150 μl of lysis buffer and transferred to a new protein LoBind tube. The beads were washed once with 2% SDS (Invitrogen, AM9820) in 50mM Tris, once with 8M Urea in 50mM Tris and then 3 times with 50mM Tris. All washing steps were performed by shaking at 800rpm at 27°Cfor 5 minutes, followed by centrifugation at 4000 x g for 2 minutes. After the final washing step, the beads were resuspended in 60 μl of 2M Urea, 50mM Tris (pH 8) containing 250 ng of trypsin (Promega, V5280) and incubated at 27°C with shaking at 800rpm for 30 minutes. Following centrifugation at 4000 x g for 2 minutes, the supernatant was collected in a fresh Eppendorf tube. Beads were then washed twice in 25 μl of 2M Urea, 50mM Tris, and the resulting supernatants were collected and pooled. To this fraction, 1 mM TECP (Promega, VB1000) was added, followed by overnight incubation at 27°C with shaking at 800rpm. The next day, trifluoroacetic acid (TFA, ThermoFisher, 85183) was added to a final concentration 0.5%, followed by centrifugation at 1400 x g. The supernatants were collected in a fresh tube and sample pH was confirmed to be at pH 3.

For peptide cleanup, C18 StageTips provided by the Functional Genomics Center Zurich (FGCZ) were prepared as follows: tips were wetted with 100% methanol (ThermoFisher 67-56-1), equilibrated with 60% acetonitrile (ACN, ThermoFisher, 75-05-8) containing 0.1% TFA and conditioned twice with 3% ACN containing 0.1%TFA. All steps were performed using 150 μl volumes, followed by centrifugation at 2000 rcf. Flow-through was discarded, and digested peptides were loaded onto the stage tips and centrifuged at 2000 rcf. Afterwards the flow-through was reapplied onto the stage tips, followed again by centrifugation at 2000 rcf. Bound peptides were washed twice with 150 μl of 3% of ACN containing 0.1% TFA and eventually eluted in 150 μl of 60% ACN containing 0.1%TFA. Eluted peptides were completely dried in a SpeedVac concentrator (ThermoFisher). Prior to the LC/MS-MS analysis, the samples were dissolved in 20 μl of 3% ACN containing 0.1% formic acid (FA, A117-05AMP) and stored at - 20°C afterwards.

### Reverse Phase Liquid Chromatography–Tandem Mass Spectrometry (LC–MS/MS)

LC/MS-MS measurements were performed on an Orbitrap Exploris 480 mass spectrometer (Thermo Fisher) coupled to an ACQUITY UPLC M-Class liquid chromatography system (Waters). Samples were loaded onto and separated on a PepSep C18 column (15 cm x 150 µm, 1.5 µm, Bruker) kept at 50 °C. For the identification of peptides in data-independent acquisition (DIA) mode, an LC method was set up to separate peptides across a 60-minute gradient at a flow rate of 300 nl/min. The gradient ranged from 95% buffer A to 5% buffer B (buffer A: 0.1% formic acid in water, buffer B: 0.1% formic acid in ACN. MS scans (400–960 m/z range, 10 m/z isolation width, 56 scan events, 33% collision energy, 800% normalized AGC target, 54 ms maximum ion time) were collected at a resolution of 30,000, and higher-energy collisional dissociation (HCD) MS/MS spectra were detected in the Orbitrap.

### Spectral library generation and protein quantification

The identification and quantification of proteins from mass spectrometry data were performed using the DIA-NN workflow^98^. The following parameters were used for the library-free search: precursor charge +2 and +3, precursor mass range 350-1500 m/z, fragment mass range 200-1800 m/z, MS1 mass accuracy 20 ppm, MS2 mass accuracy 20 ppm, trypsin/P enzyme specificity allowing two missed cleavages, cysteine carbamidomethylation as a fixed modification, and N-terminal methionine excision as a variable modification. The in-silico spectra were generated using a canonical human protein database (Uniprot ID: UP000005640), concatenated with common protein contaminants, with the maximum false discovery rate (FDR) set to 0.01.

### Statistical modeling of the proteomics data

The R packages *prolfqua* and *prolfquapp* were used to analyze differential expression with respect to group differences^99, 100^, including the estimation of confidence intervals and false discovery rates for all quantifiable proteins. From the output report.tsv file generated by DIA-NN, that contains the precursor ion abundances for each raw file, protein abundances were determined by first aggregating precursor abundances to peptidoform abundances, followed by estimation via Tukey’s Median Polish. Protein abundances were transformed using robscale before fitting the linear models. Under the same protein group, only the one with the highest numbers of peptides identified was kept. PCA and volcano plots illustrating group differences and differentially expressed proteins were generated using the R package *ggplot2*^101^. Differentially expressed proteins were filtered for statistical significance (FDR < 0.05) and a group difference greater than 1. The mass spectrometry proteomics data have been deposited to the ProteomeXchange Consortium via the PRIDE^102^ partner repository with the dataset identifier PXD071669 (for reviewer account access; pw-H7tDZa5GkpFd) for nuclear and cytoplasmic TDP-43 interactors, and PXD071675 (for reviewer account access; pw-WwJI4pQQMLHS) for the patient seeding experiment.

### Functional enrichment analysis

Functional enrichment analysis was performed on genes within significant modules using the *clusterProfiler* R package (v4.16.0)^103^. Gene Ontology (GO) enrichment analysis was conducted on Biological Processes (BP), Molecular Function (MF) and Cellular Compartments (CC) employing the *enrichGO* function. Gene symbols were mapped to Entrez IDs using the org.Hs.eg.db annotation database. Enrichment significance was determined based on an adjusted p-value threshold of 0.05. KEGG pathway enrichment analysis was performed using the *enrichKEGG* function, specifying the human database (hsa). Results were visualized using dot plots and bar plots, displaying enriched terms, significance levels, and gene ratios.

### Overlap analysis of the differential interactome of TDP-43

To identify shared and unique TDP-43 differential interactors across experimental comparisons and external datasets, pairwise overlap analyses were performed. Published proteomics datasets associated with TDP-43 pathology were collected from ten studies or databases^52–61^. Additionally, TDP-43 protein-protein interaction data were obtained from BioGRID database^59^. The Jaccard similarity of the comparisons was calculated and the significance of overlaps was assessed using Fisher’s Exact Test. Genes of non-human species were converted to human orthologs for comparison using the *biomaRt* package (v3.21), while human datasets were processed directly or updated to reflect current gene nomenclature. For the full list of genes and references for the datasets analyzed, see supplementary document 1.

### Statistical analyses

To explain the variation in the seeding capacity of the patient-derived materials, which can be attributed to variation in the collected variables about these individuals, we applied multiple linear regression to determine whether some of these explanatory variables about the individuals have a linear relationship with the response, and to identify which subsets of the variables may contain non-redundant information about the response. We set the amount of neo-aggregate formation at 10 d.p.i. in the WT cell line, and the percentage of mScarlet-positive cells at 18 d.p.i. as the response variables to avoid redundancy, since the result on both timepoints were highly correlated with a Pearson’s correlation coefficient of 0.99. In addition, we investigated whether accounting for clinical factors clinical factors including postmortem delay (PMD), age at death, and sex would improve the model for explaining the seeding effect. Additionally, multiple variables from biochemical characterization appeared to be highly correlated with each other. For these reasons, we initiated a full model containing all non-redundant variables collected, with backward stepwise selection to identify the optimal subset of predictors to avoid overfitting and to improve model interpretability. The modeling was performed by implementing the *LinearRegression* class from *scikit-learn* machine learning library. Backward stepwise selection was employed to identify the optimal subset of predictive features for the linear regression model by iteratively removed features one at a time, retaining the subset that maximized the coefficient of determination (R²) at each step. Given the limited sample size relative to the number of potential predictors, we monitored for overfitting throughout the selection process by examining the discrepancy between training R² and cross-validated R² for each candidate model. Cross-validation was performed using 5-fold cross-validation, with a minimum threshold of 3 observations per feature in each fold to ensure stable model evaluation. Model performance was evaluated across all possible feature subsets by Bayesian Information Criterion (BIC) which penalizes model complexity especially for small sample sizes. For each candidate model, we calculated standard errors, t-statistics, p-values, and 95% confidence intervals for all regression coefficients using ordinary least squares estimation. Statistical significance was assessed at the α = 0.05 level.

Other statistical analyses were performed with Prism software (GraphPad, v10.5.10.) unless otherwise specified. Values are displayed as individual data points with mean and standard deviation (SD). Statistical significance is indicated as follows: p < 0.05 (*), p < 0.01 (**), p < 0.001 (***), p < 0.0001 (****).

---

**Supplementary Table 1.** Overlap of TDP-43 interactors with external datasets. Numbers of shared and unique differential TDP-43 interactors across experimental conditions in this work and 10 other external datasets were compared using absolute overlap counts.

| Dataset | Model | Cheng_WT | Cheng_dNLS | Blokhuis | Helmold | Freibaum | Riemenschneider | Evangelista | Feneberg | Chou | Schreiber | BioGRID | Zhong_WT | Zhong_mNLS |
| --- | --- | --- | --- | --- | --- | --- | --- | --- | --- | --- | --- | --- | --- | --- |
| Cheng_WT | HEK293 | 314 | 26 | 38 | 24 | 65 | 57 | 210 | 69 | 51 | 50 | 79 | 35 | 26 |
| Cheng_dNLS | HEK293 | 26 | 179 | 14 | 10 | 24 | 23 | 75 | 21 | 23 | 20 | 32 | 18 | 17 |
| Blokhuis | NSC-34 | 38 | 14 | 132 | 36 | 81 | 22 | 91 | 32 | 33 | 56 | 80 | 62 | 21 |
| Helmold | literature review | 24 | 10 | 36 | 155 | 105 | 18 | 78 | 26 | 14 | 28 | 125 | 29 | 18 |
| Freibaum | HEK293 | 65 | 24 | 81 | 105 | 253 | 40 | 170 | 55 | 39 | 64 | 220 | 64 | 43 |
| Riemenschneider | rat primary neuron | 57 | 23 | 22 | 18 | 40 | 200 | 97 | 39 | 23 | 34 | 49 | 17 | 18 |
| Evangelista | HEK293A | 210 | 75 | 91 | 78 | 170 | 97 | 1896 | 109 | 134 | 251 | 236 | 199 | 96 |
| Feneberg | Mouse ES-cell derived motor neurons | 69 | 21 | 32 | 26 | 55 | 39 | 109 | 270 | 26 | 31 | 66 | 24 | 13 |
| Chou | N2A | 51 | 23 | 33 | 14 | 39 | 23 | 134 | 26 | 219 | 59 | 46 | 60 | 16 |
| Schreiber | HEK293 | 50 | 20 | 56 | 28 | 64 | 34 | 251 | 31 | 59 | 769 | 100 | 365 | 2 |
| BioGRID | literature review | 79 | 32 | 80 | 125 | 220 | 49 | 236 | 66 | 46 | 100 | 560 | 91 | 46 |
| Zhong_WT | SH-SY5Y | 35 | 18 | 62 | 29 | 64 | 17 | 199 | 24 | 60 | 365 | 91 | 713 | 10 |
| Zhong_mNLS | SH-SY5Y | 26 | 17 | 21 | 18 | 43 | 18 | 96 | 13 | 16 | 2 | 46 | 10 | 473 |

**Supplementary Table 2.**
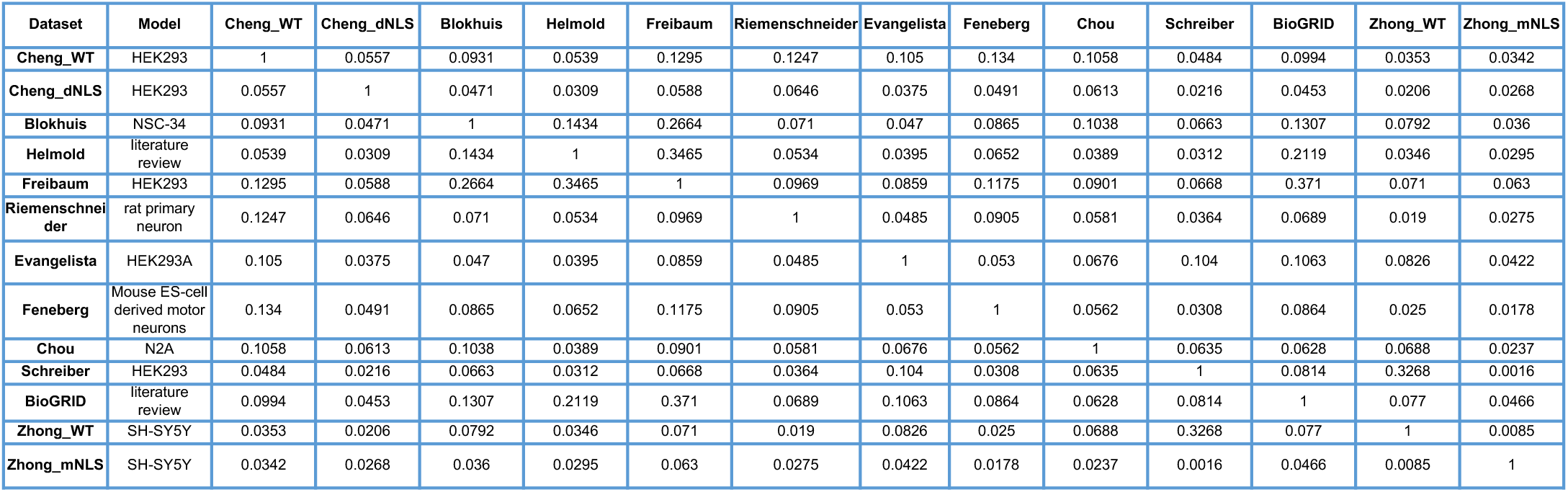
Similarity of TDP-43 interactors with external datasets. Comparison of differential TDP-43 interactors across experimental conditions in this work and 10 other published datasets using Jaccard similarity coefficients (intersection/union).

**Supplementary Table 3.**
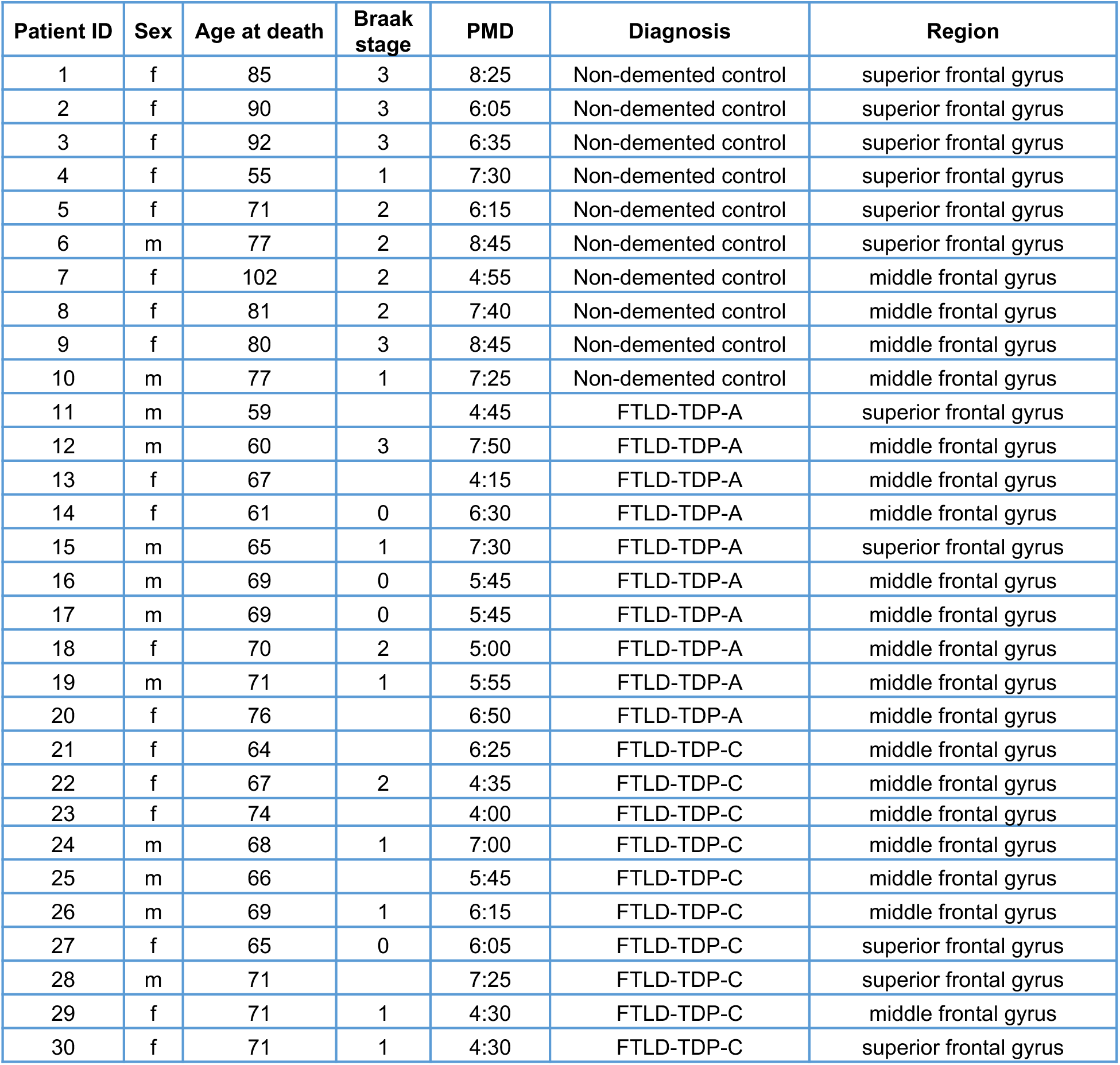
Patient cohort information. Patient information of the postmortem samples used in this study, which were provided by the Netherlands Brain Bank (NBB), Netherlands Institute for Neuroscience, Amsterdam.

**Supplementary Figure 1.**
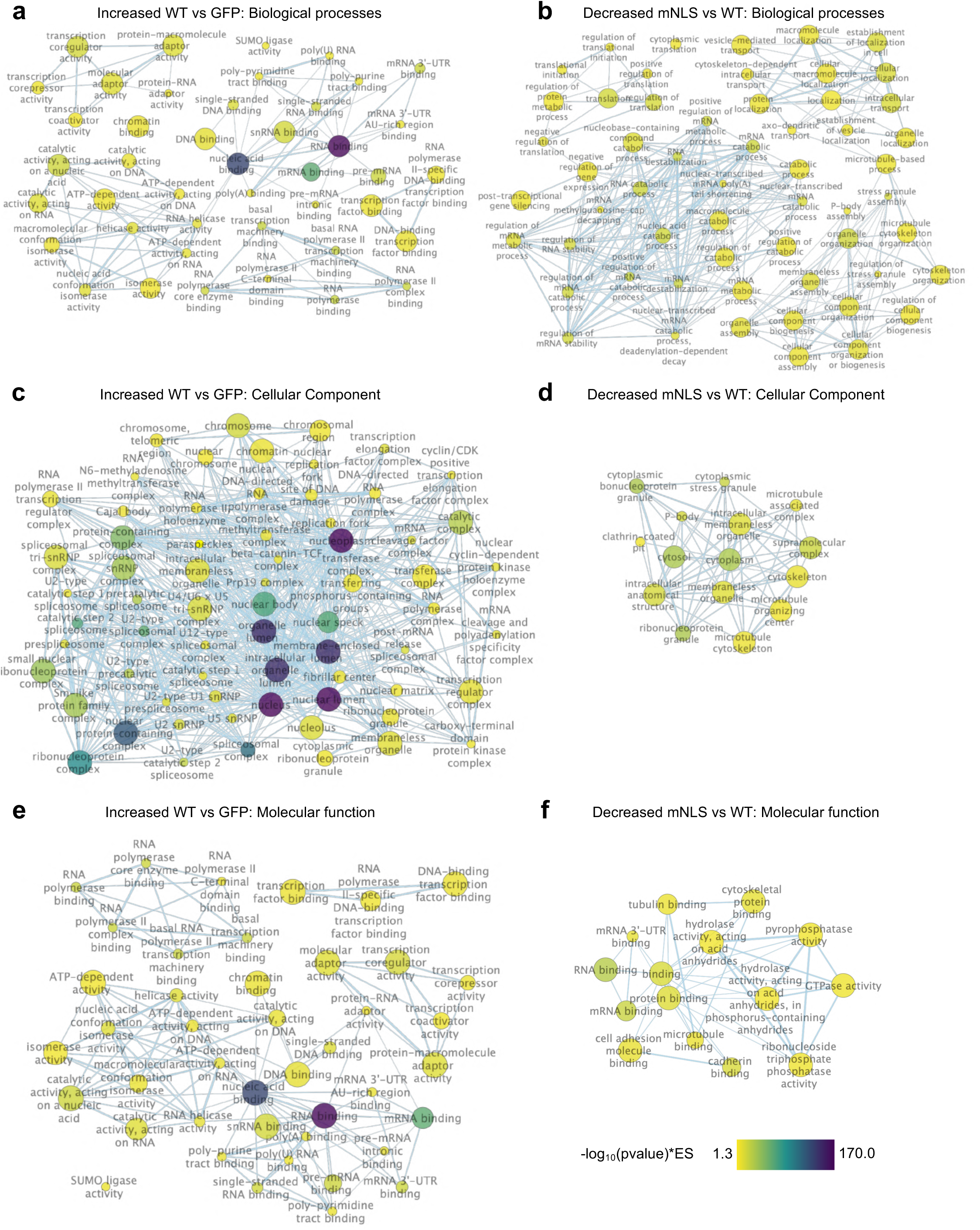
Enrichment analysis of TDP-43 nuclear and cytoplasmic interactors. Enriched gene ontology (GO) terms for increased interactions in the comparison of WT vs. GFP and decreased for WT vs. mNLS visualized in the *Enrichment Map* application (v. 3.5.0) in *Cytoscape* (v. 3.10.3), filtered for -log_10_(adjusted p-values) significance based on normalized enrichment score (NES) between 1.3 and 170 inclusive. Nodes represent gene sets, and node size represents the number of genes in each set, whereas edges represent overlap (similarity) between gene sets. Edge width shows the number of overlapping genes and node fill shows -log_10_(adjusted p-values) significance (NES).

**Supplementary Figure 2.**
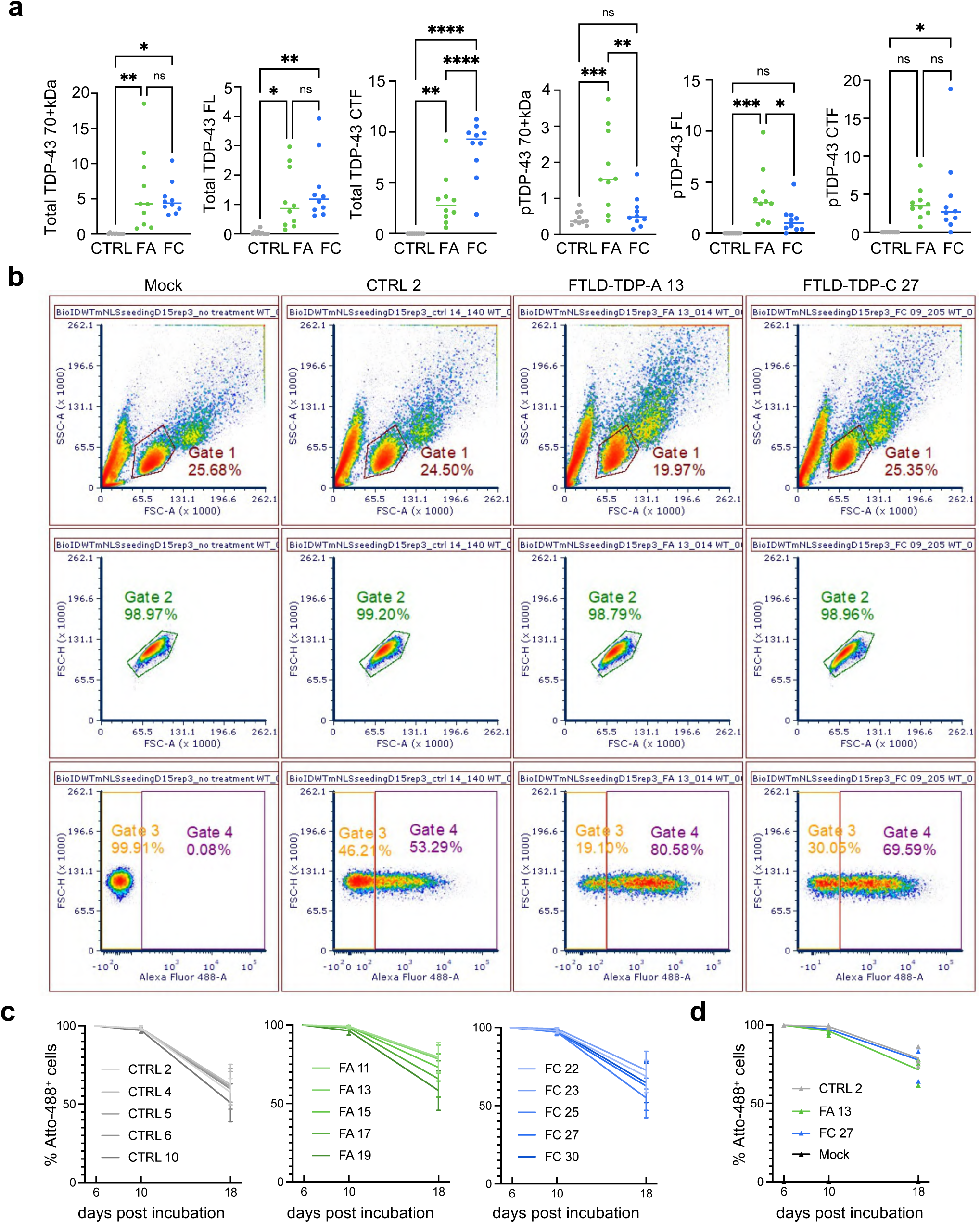
BioID-SH-SY5Y cells readily internalized patient-derived seeds. (**a**) Additional quantifications of the western blots were performed based on the intensity for various TDP-43 species observed in SarkoSpin pellets: hyperphosphorylated/polyubiquitination smear at 70 kDa and above (70+kDa), TDP-43 full length (FL) at ∼45 kDa, and the C-terminal fragments of ∼25 kDa (CTFs) for total TDP-43 and pTDP-43. Statistical analyses performed using one-way ANOVA with Tukey’s multiple comparisons (*p < 0.05, **p < 0.01, ***p < 0.001, ***p < 0.0001, ns = not significant). Y-axis values represent fluorescence intensity expressed in arbitrary units (A.U.). (**b**) Representative FACS scatter plots of SH-SY5Y cells stably expressing BioID2-TDP-43WT at 18 d.p.i. with Atto-488-labeled patient-seeds (SKS-488), *n*=3. (**c**) Percentage of BioID2-TDP-43WT cells with internalized TDP-43 aggregates measured by FACS after incubation with labeled patient-seeds with 5 individuals per condition as biological replicates from non-degenerative control (CTRL, gray), FTLD-TDP-A (FA, green), FTLD-TDP-C (FC, blue). (**d**) Percentage of BioID2-TDP-43mNLS cells with internalized TDP-43 aggregates, measured by FACS after incubation with labeled patient-seeds with a subset of individuals one for each treatment condition, as well as one non-seeding, mock control (black). Lines indicate the mean of three independent experimental repetitions and error bars the standard deviation.

**Supplementary Figure 3.**
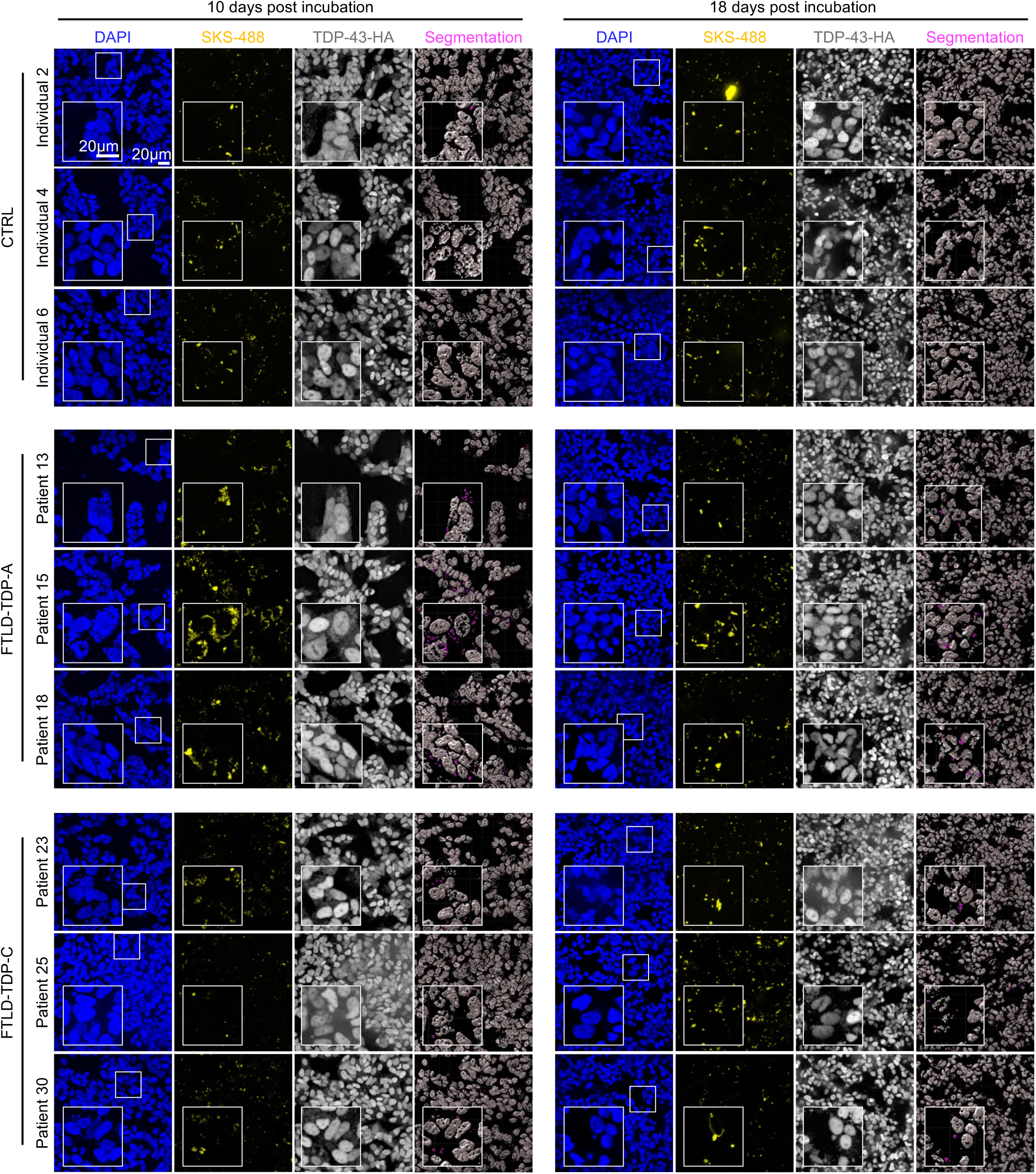
Patient-derived seeds induce the aggregation of endogenous TDP-43. Representative confocal immunofluorescence images of seeded BioID2-TDP-43WT cells taken at on 10 and 18 d.p.i. 3D reconstructed objects of endogenous TDP-43-HA signal, displayed in white, generated with automatic segmentation in Imaris through intensity thresholding and splitting touching objects. Neoaggregates are highlighted in magenta and depict TDP-43-HA colocalizing with exogenous fluorescently labeled aggregates with an overlapping volume of over 30%. Scale bar: 20 μm.

**Supplementary Figure 4.**
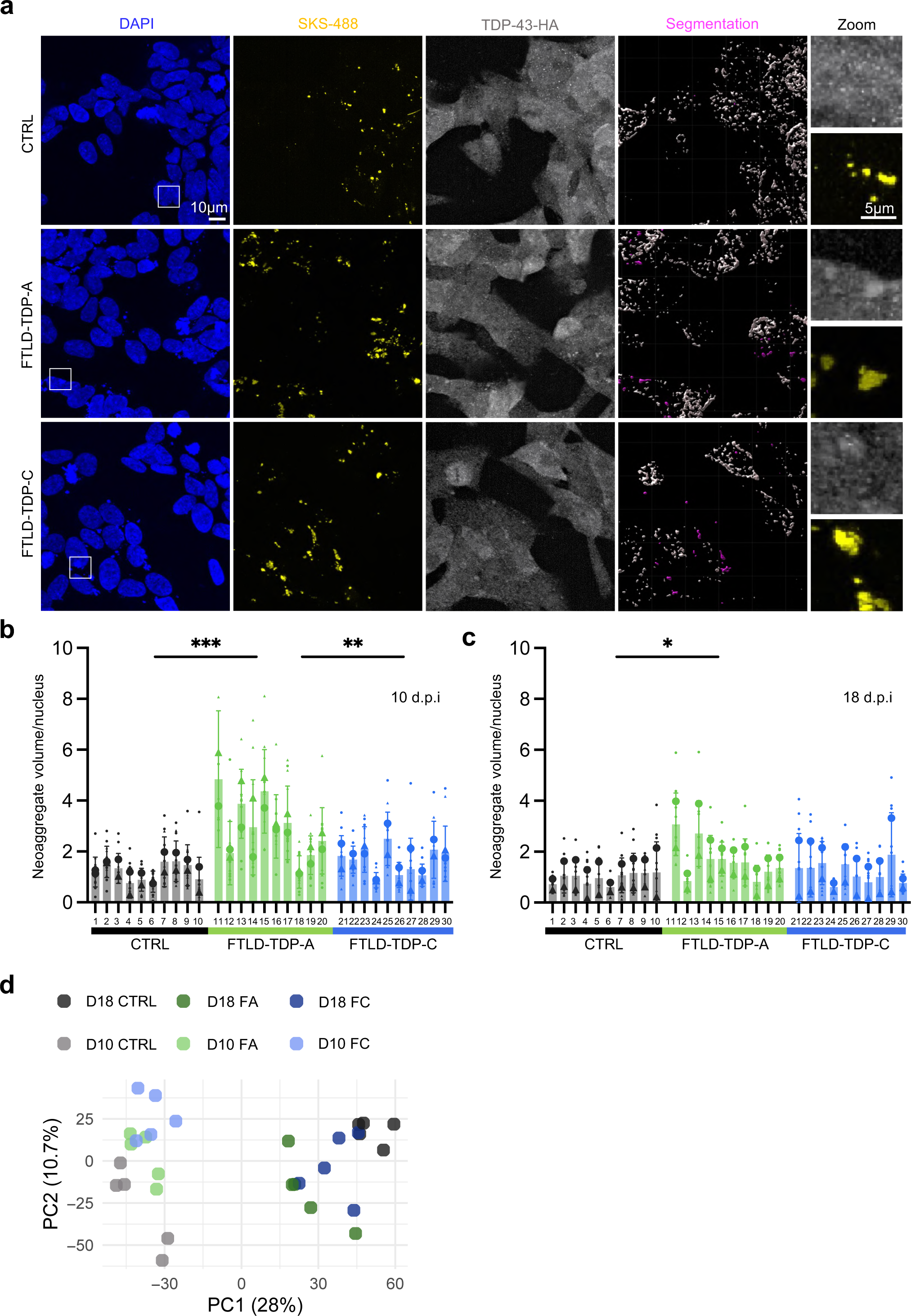
Patient seeding induces aggregation and proteomic changes in BioID2-TDP-43mNLS cells. (**a**) Representative confocal immunofluorescence images of seeded BioID2-TDP-43mNLS cells at 10 and 18 d.p.i. 3D reconstructed objects of endogenous TDP-43-HA (white) generated with automatic segmentation in Imaris through intensity thresholding and splitting touching objects. Neoaggregates (magenta) depict TDP-43-HA colocalizing with exogenous fluorescently labeled aggregates with an overlapping volume of over 30%. Scale bar: 10 μm and 5 μm in zoomed images. (**b, c**) Image quantification of the neoaggregate volume, normalized to the count of cells per field-of-view (FOV) in BioID2-TDP-43mNLS cells at (**b**) 10 and (**c**) 18 d.p.i. Smaller dots represent FOVs, and larger dots represent the mean of values from two independent experiments, with 10 biological replicates (individuals) per group (n=10). Error bars show the standard deviation. Statistical analyses for the comparisons between CTRL, FTLD-TDP-A and FTLD-TDP-C using a one-way ANOVA with Tukey’s multiple comparisons (*p < 0.05, **p < 0.01, ***p < 0.001). (**d**) Principal component analysis of proteomics data demonstrating separation of the experimental groups (CTRL, FTLD-TDP-A, FTLD-TDP-C). Each dot represents sample treated with seeds from one

**Supplementary Figure 5.**
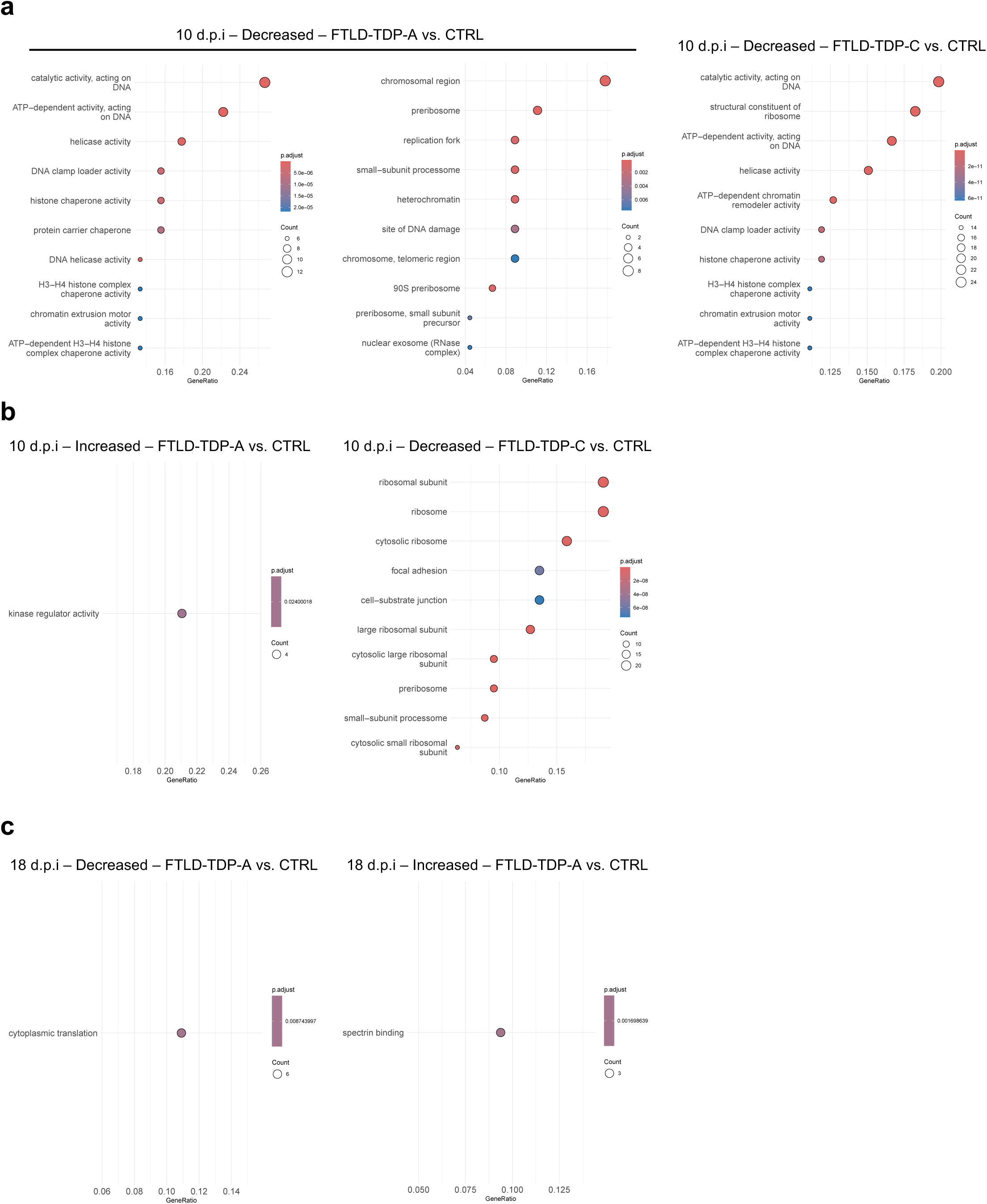
Differentially enriched subtype-specific TDP-43 interactors. (**a-b**) Differentially enriched TDP-43 interactors across the following comparisons at 10 d.p.i.: decreased FTLD-TDP-A and FTLD-TDP-C vs. CTRL, and (**b**) increased FTLD-TDP-A vs. CTRL and decreased FTLD-TDP-C vs. CTRL, and (**c**) at 18 d.p.i. decreased and increased for FTLD-TDP-A vs. CTRL. Dot sizes represent the number of genes in each pathway (count), while dot colors indicate pathway significance based on adjusted p-values (p.adjust). The GeneRatio on the x-axis represents the proportion of the differential interactors associated with each pathway.

**Supplementary Figure 6.**
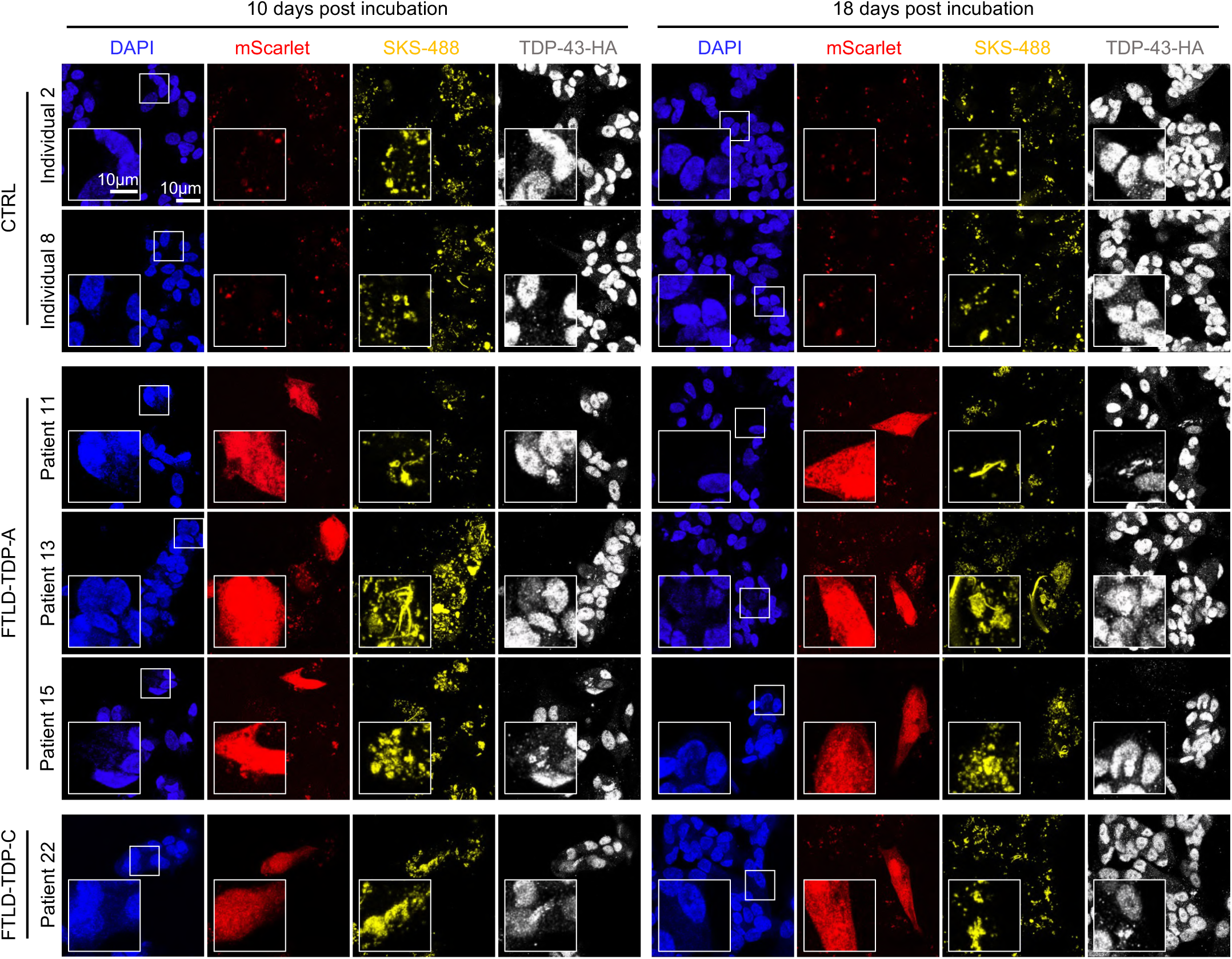
Seeded aggregation with patient material induces TDP-43 loss of function. Representative confocal immunofluorescence images of seeded TDP-REG cells taken at on 10 and 18 d.p.i. Scale bar: 10 μm. Representative immunofluorescence images taken on 18-days p.i. Dotted lines delineate the outline of mScarlet-positive (mScarlet+) cells, whereas dashed lines mark the nuclear region of the same cell that displayed reduced endogenous TDP-43 intensity. The white arrowhead remarks an example of internalized filamentous FTD-A aggregate with the recruitment of endogenous TDP-43 in a mScarlet-positive cell. Scale bar: 5 μm.

**Supplementary Figure 7.**
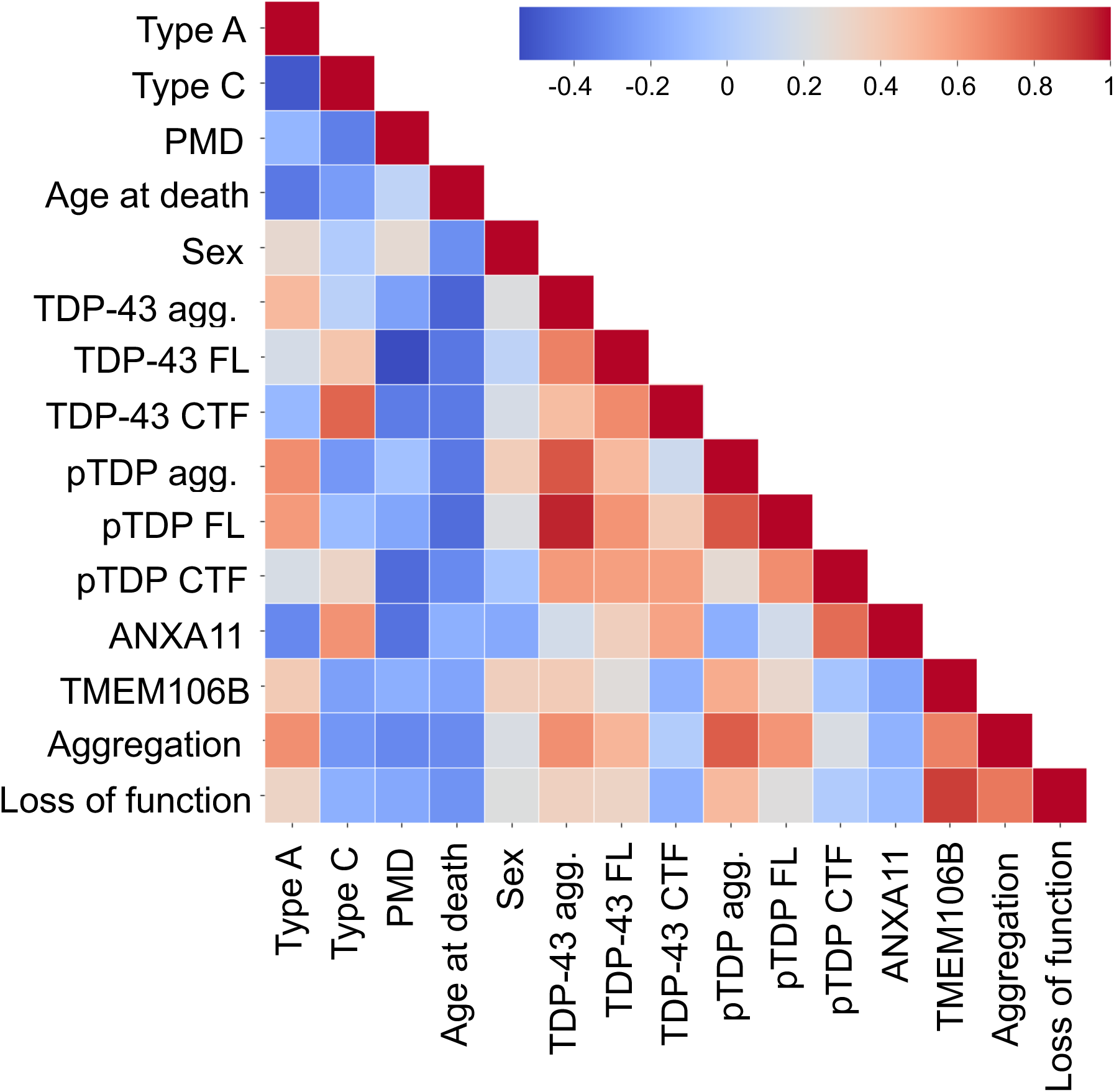
Associations of biochemical and clinical factors and seeding outcome. Pairwise correlations between the clinical and biochemical variables, characterized by TDP-43 species separated by molecular weights on western blots, regarding the individuals, with the capacity of inducing neo-aggregate formation of the seeds derived from those individuals were assessed using Pearson’s correlation coefficient. Results were visualized as a correlation heatmap with the color scale representing correlation strength and direction.

**Supplementary Figure 8.**
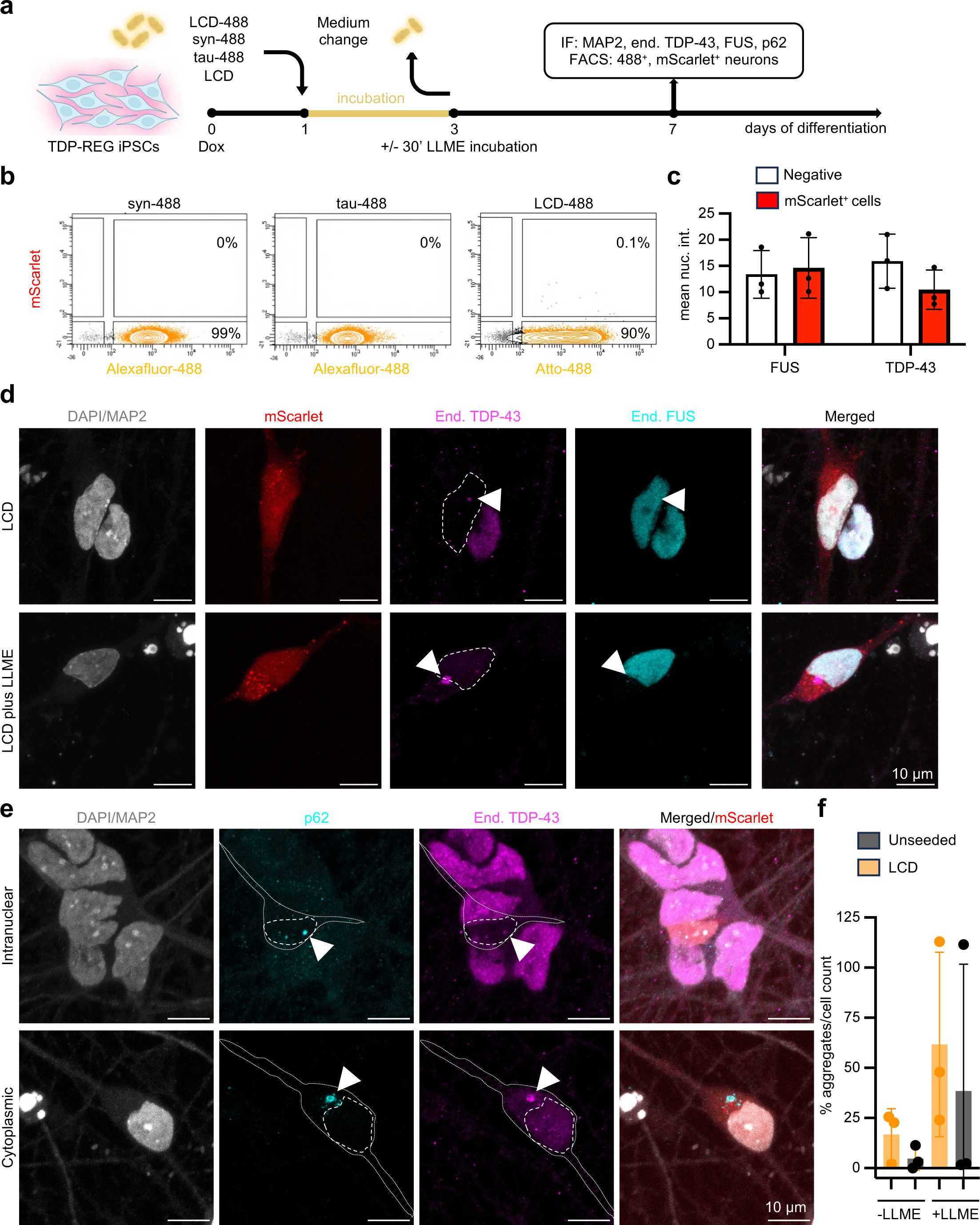
Lysosomal escape unlocks TDP-43 seeding in human neurons. (**a**) Schematic representation of our seeding model using TDP-REG *NGN2* iPSCs. One day after plating, neuronal precursor cells (NPCs) are incubated with 200 nM of unlabeled or fluorescently labeled TDP-43 LCD (LCD; LCD-488), or a-synuclein or tau fibrils. Two days after fibril incubation, NPCs are treated for 30 min with 1 mM L-leucyl-L-leucine methyl ester (LLME) (where specified), after which the medium is completely replaced. NPCs undergo differentiation until day 7, when they are fixed for immunofluorescence or collected for FACS. (**b**) FACS contour plots show the populations of Alexafluor-488-, Atto-488- and mScarlet-positive neurons after seeding with 200 nM of LCD, a-synuclein, or tau K18 labeled fibrils (n=3). (**c**) Quantification of the mean nuclear intensity of endogenous TDP-43 and FUS in mScarlet-positive neurons upon seeding with LCD combined with LLME treatment, based on images acquired at differentiation day 7 (n=3; 10 FOVs analyzed per replicate containing at least one mScarlet-positive neuron). Each single dot represents the mean nuclear intensity of mScarlet-positive neurons for each of the three independent experiments. Error bars show standard deviation (SD). (**d**) Representative immunofluorescence images taken at differentiation day 7 show endogenous TDP-43 (magenta) and FUS (cyan) in mScarlet-positive neurons with intranuclear (upper panel) or cytoplasmic (lower panel) TDP-43 pathology after seeding with 200 nM LCD fibrils. Dotted lines highlight the nuclear profile overlayed on the TDP-43 signal, which is reduced in the same nucleus where FUS signal is maintained. Scale bar 10 µm (n=3). (**e**) Representative immunofluorescence images taken at differentiation day 7 show mScarlet-positive neurons after seeding with 200 nM of unlabeled LCD displaying two different types of inclusions positive for endogenous TDP-43 (magenta) and p62 (cyan): intranuclear (top panels) and cytoplasmic (lower panels). Dotted lines show the body of the mScarlet-positive neurons and highlight the nuclear profile overlayed on the p62 and TDP-43 signal. Arrowheads mark p62 signal co-localizing with endogenous TDP-43. Scale bar 10 µm (n=3). (**f**) Image-based quantification of p62 and TDP-43 aggregates expressed as a percentage of the number of counted cells (n=3). Each single dot represents the mean percentage of aggregates over total cell count for each independent replicate. Error bars show SD.

**Supplementary Figure 9.**
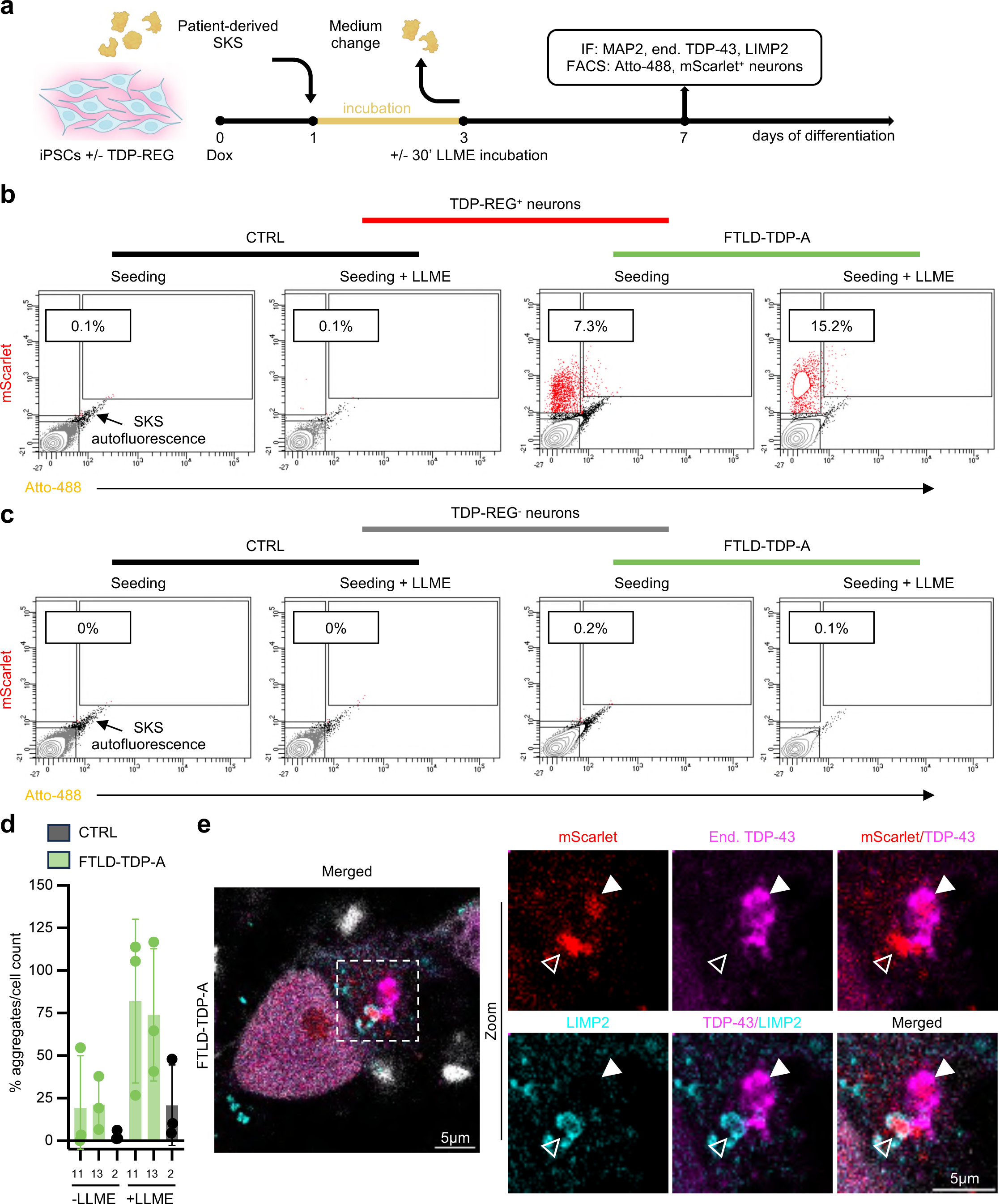
TMEM106B promotes TDP-43 seeding with patient-derived TDP-43 fibrils. (**a**) Schematic representation of our seeding model using TDP-REG and non-TDP-REG *NGN2* iPSCs. One day after plating, neuronal precursor cells (NPCs) are incubated with patient-derived SarkoSpin extracts (SKS) (5.3% of the total resuspended and sonicated pellet per cm^2^ well surface area). Two days after fibril incubation, NPCs are treated for 30 min with 1 mM L-leucyl-L-leucine methyl ester (LLME) (where specified), followed by full medium change. NPCs undergo differentiation until day 7, when they are fixed for immunofluorescence or collected for FACS. (**b**) Representative FACS contour plots show mScarlet and Atto-488 signal of TDP-REG *NGN2* neurons seeded with SKS patient-derived material, in the absence or presence of LLME, and (**c**) analogous plots obtained by seeding *NGN2* neurons devoid of the TDP-REG system. (**d**) Image-based quantification of p62 and TDP-43 aggregates expressed as a percentage of the number of counted cells (n=3). Each single dot represents the mean percentage of aggregates over total cell count for each independent replicate for one CTRL (nr.2) and two FTLD-TDP-A (nr.11 and 13). Error bars show standard deviation (SD). (**e**) Immunofluorescence images taken at differentiation day 7 show one mScarlet-positive neuron seeded with FTLD-TDP-A SKS (nr.15) plus LLME, as a representative example of how recruiting aggregates were quantified. LIMP2 staining (cyan) shows lysosomal sequestration of exogenous SKS material (red, autofluorescent in the 568 channel used for mScarlet). Exogenous SKS material was considered a recruiter if colocalizing with endogenous TDP-43 (magenta). White arrowheads mark a recruiting aggregate, while the empty ones mark a non-recruiter. Scale bar 5 µm (n=3)

